# Towns and Trails Drive Carnivore Connectivity using a Step Selection Approach

**DOI:** 10.1101/2021.02.24.432739

**Authors:** Jesse Whittington, Robin Baron, Mark Hebblewhite, Adam T. Ford, John Paczkowski

**Affiliations:** Park Canada, Banff National Park Resource Conservation. PO Box 900, Banff, Alberta, Canada. T1L 1K2; Wildlife Biology Program, Department of Ecosystem and Conservation Sciences, W.A. Franke College of Forestry and Conservation, University of Montana, 32 Campus Drive, Missoula, MT, USA.59801; Department of Biology, Faculty of Science, University of British Columbia, Kelowna, BC, Canada. V1V 1V7; Alberta Environment and Parks, Kananaskis Region, 201, 800 Railway Avenue, Canmore, Alberta, Canada. T1W 1P1

**Keywords:** connectivity, conservation, corridors, movement ecology, human development, resource selection, step selection, utilization distribution

## Abstract

Global increases in human activity threaten connectivity of animal populations. Protection and restoration of animal movement corridors requires robust models to forecast the effects of human activity on connectivity. Recent advances in the field of animal movement ecology and step selection functions offer new approaches for estimating connectivity. We show how a combination of hidden Markov movement models and step selection functions can be used to simulate realistic movement paths with multiple behavioral states. Simulated paths can be used to generate utilization distributions and estimate changes in connectivity for multiple land use scenarios. We applied movement models to 20 years of grizzly bear (*Ursus arctos*) and gray wolf (*Canis lupus*) data collected in and around Banff National Park, Canada. These carnivores avoided areas near towns in all seasons, avoided areas of high trail density in most seasons, and campgrounds during summer and fall. We simulated movement paths for three landscape scenarios: reference conditions with no anthropogenic development, current conditions, and future conditions with expanded town footprints and trail networks. We counted the number of paths that crossed valley-wide, digital transects through mountain tourist towns of Banff and Canmore, Alberta. We divided current and future crossing rates by the reference crossing rates to estimate connectivity. Current connectivity rates ranged between 7 and 45% of reference values with an average of 21% for grizzly bears and 25% for wolves. Potential town expansion and increased development of trails further decreased connectivity an average of 6% in future scenarios. Anthropogenic developments reduced the amount of available high quality large carnivore habitat in the Bow Valley by an average of 14% under current conditions and 16% under future conditions. Our approach for estimating connectivity provides a robust and flexible method for combining movement models with step selection analyses to estimate connectivity for a variety of species.

## Introduction

Global increases in human activity threatens wildlife populations and as a result, many conservation programs have increased their focus on ecological connectivity (Hilty et al. 2020). Connectivity analyses of animal movement are frequently used to identify likely dispersal routes between populations (Fattebert et al. 2015, Zeller et al. 2018), seasonal migrations routes (e.g. Fullman et al. 2021), and to highlight natural and anthropogenic pinch points to movement (i.e. wildlife corridors) as priority areas for conservation (Chetkiewicz and Boyce 2009, Suraci et al. 2020). Within an animal’s home range, wildlife corridors facilitate movements important for reproduction, accessing seasonal resources, and predator-prey processes (Hebblewhite 2005, Panzacchi et al. 2016). At broader scales, connectivity facilitates dispersal (Benz et al. 2016), gene flow, and demographic rescue of subpopulations (Marrotte et al. 2017, Lamb et al. 2020). A wide variety of approaches have been used to estimate connectivity of animal movements, with varying degrees of success (Calabrese and Fagan 2004, Zeller et al. 2018). Emerging techniques in the field of movement ecology offer new opportunities to develop stronger links between movement behavior and estimates of connectivity (e.g. Hooten et al. 2020).

Movement models and step selection analyses offer a complementary approach for estimating connectivity either from model predictions (Buderman et al. 2018, Hooten et al. 2020) or from simulated paths (Palmer et al. 2011, Quaglietta and Porto 2019, Zeller et al. 2020). Step selection analyses are a subset of spatial point-process models that are increasingly used to estimate relative selection of resources (Fortin et al. 2005), to understand the effects of human activity on animal movement behaviour (e.g. Suraci et al. 2019), and to create utilization distributions (UDs) that predict spatial variation in intensity of habitat use (Signer et al. 2017).

Step selection analyses have become increasingly accessible for practitioners through the development of statistical packages in R (Avgar et al. 2016, Signer et al. 2019, Muff et al. 2020). Several studies have incorporated step selection functions (SSFs) into connectivity analyses by first creating spatial predictions of habitat use and then transforming predictions into resistance layers for cost-distance or circuit theory analyses (Zeller et al. 2018, Brennan et al. 2020, Suraci et al. 2020). Others have used the derived resistance surfaces to simulate animal movements (Quaglietta and Porto 2019, Jayadevan et al. 2020, Zeller et al. 2020). For example, Merkle et al. (2019) simulated movements directly from an SSF to forecast migration routes. Simulated individual-based paths are appealing because they can incorporate sequential, probabilistic movement decisions related to landscape features, speed of travel, and directional persistence (Avgar et al. 2016). Moreover, simulating animal movements is considered the best practice for generate unbiased UDs from SSFs (Signer et al. 2017). Simulations, while computational intensive, can easily be applied to multiple land use scenarios. Movement simulations from SSFs offer a promising method for assessing the cumulative effects of multiple landscape features on animal movement paths, intensity of use, and connectivity.

Realistic simulations need to accommodate the underlying factors that influence animal movement including seasonal (Zeller et al. 2019, Brennan et al. 2020, Zeller et al. 2020) and temporal (Gaynor et al. 2018, Lamb et al. 2020) variability in resource selection and state-specific movement behaviors (Michelot et al. 2016). For example, animals often have low directional persistence and low speed of travel when feeding and resting in slow states and have strong directional persistence and higher speed of travel when travelling in fast states (Fryxell et al. 2008). Such behavioral states are overlooked in classical circuit theory and cost-distance type connectivity models. And failure to incorporate behavioral state into SSFs can lead to biased UDs, poor estimates of connectivity, and misidentification of wildlife corridors (Abrahms et al. 2017). Finally, responses to human activity and estimates of connectivity can vary widely among species (e.g. Rogala et al. 2011, Brennan et al. 2020, Nickel et al. 2020). From a conservation perspective, focussing on the most sensitive species should increase connectivity for most other wildlife (Meurant et al. 2018, Lamb et al. 2020).

Large carnivores are an important consideration for landscape-scale measures of connectivity for a number of reasons. First, these iconic and charismatic species are often selected as conservation ‘flagship’ and umbrella species, meaning they hold a particularly deep value for the public and management agencies (Ray et al. 2013). Second, the potential threat of carnivores to human safety requires a detailed understanding of how animals move through human-dominated landscapes (Buchholtz et al. 2020, Lamb et al. 2020). Third, large carnivores have the potential to affect community-level processes through top-down control on prey abundance and thus trophic cascades (Hebblewhite et al. 2005, Hebblewhite and Merrill 2011, Ripple et al. 2014). Consequently, understanding how movements of carnivores are affected by connectivity and corridor design policies (Ford et al. 2020) is an important step towards better management of ecosystem-level process (Terborgh et al. 1999).

The novel approaches to quantifying connectivity afforded through SSF-derived simulations may support better land use decision making for carnivore conservation. Here, we focused on assessing carnivore connectivity in a transboundary region of Banff National Park (BNP), AB, Canada where transportation infrastructure, outdoor recreation, and urban areas occupy much of the prime habitat in the valley bottoms of the mountainous landscape. We focus on the movements of grizzly bears (*Ursus arctos*) and wolves (*Canis lupus*) because of their management relevance, threatened status, and important ecological roles (e.g. Hebblewhite et al. 2005). We used 20 years of grizzly bear and wolf telemetry data to develop seasonal hidden Markov models and SSFs. We simulated animal paths from the movement models and SSFs to assess changes in UDs and connectivity. Based on grizzly bear and wolf responses to human activity in other studies (Whittington et al. 2005, Hebblewhite and Merrill 2008, Lamb et al. 2020), we expected grizzly bears and wolves to select linear features as efficient travel routes while avoiding areas near towns and areas with high trail density. We expected avoidance to be most pronounced and connectivity to be lowest during peak tourist visitation in summer. Finally, we expected that connectivity around towns would decrease from current to future conditions due to an expanded town footprint and increased recreational trail density (Gutzwiller et al. 2017). Building on the growing field of movement ecology, we provide a flexible approach to generate movement-based estimates of connectivity that can be applied to other taxa and systems.

## Materials and Methods

### Study area

The study area encompassed 17,450 km^2^ of the Canadian Rockies within and adjacent to BNP (51.2° N, 115.5° W, Appendix S1: Figure S1). We defined the extent of the study area based on movements of radio-collared wolves and grizzly bears monitored from 2000 to 2020. The study area contained rugged topography, short summers and long cold winters. See Whittington et al. (2019) for a description of vegetation and the predator-prey community.

The study area contained the tourist towns of Banff and Canmore and several hamlets that occupied the centre of the Bow Valley. Linear features such as the Trans Canada highway, a national railway, and secondary roads bisected the study area. Like many global protected areas (Wittemyer et al. 2008), human activity within the study area increased steadily over the last 20 years (Alberta Environment and Parks 2018), with the potential for increasing impacts on wildlife connectivity (Gutzwiller et al. 2017). BNP currently receives over 4 million visitors per year, mostly concentrated in summer. Most anthropogenic developments and recreational activities were concentrated near roads within the Bow Valley. Backcountry areas in the northeastern portion of the study area received minimal human use.

### Telemetry data

Researchers fit wolves and grizzly bears with Global Positioning System (GPS) collars to collect data from 2000 to 2020. Researchers captured and collared grizzly bears using a combination of culvert traps and free-range darting and wolves using a net shot from a helicopter under University and Federal government capture and Animal Care permits (see Appendix S1 for summary of permits). Researchers programmed most collars to collect GPS locations every two hours. GPS collars had high fix rates with low habitat-induced fix-rate bias (Hebblewhite et al. 2007). We obtained a large sample of locations from both front and backcountry areas (Figure 2, Appendix S1: Figures S1 – S3).

**Figure 1.**
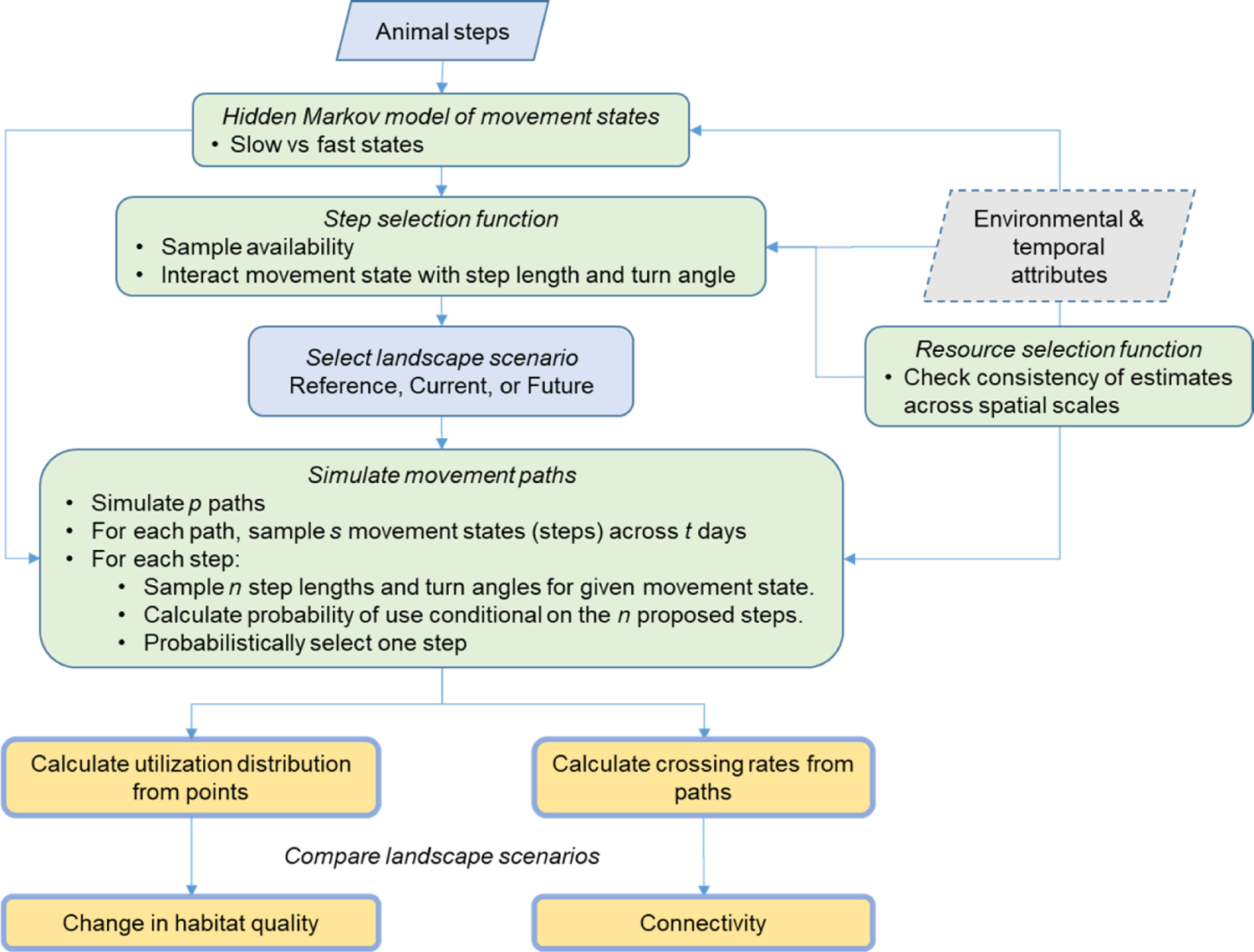
Workflow to assess connectivity and change in habitat quality using hidden Markov models, SSFs, and RSFs. We classified movement behaviors into slow and fast states and then used those states in SSF models and in path simulations. Simulated points can be used to estimate UDs and changes in habitat quality. Connectivity can be measured by comparing movement rates through corridors, across transects, or between patches relative to a reference model of movement with no anthropogenic development.

**Figure 2.**
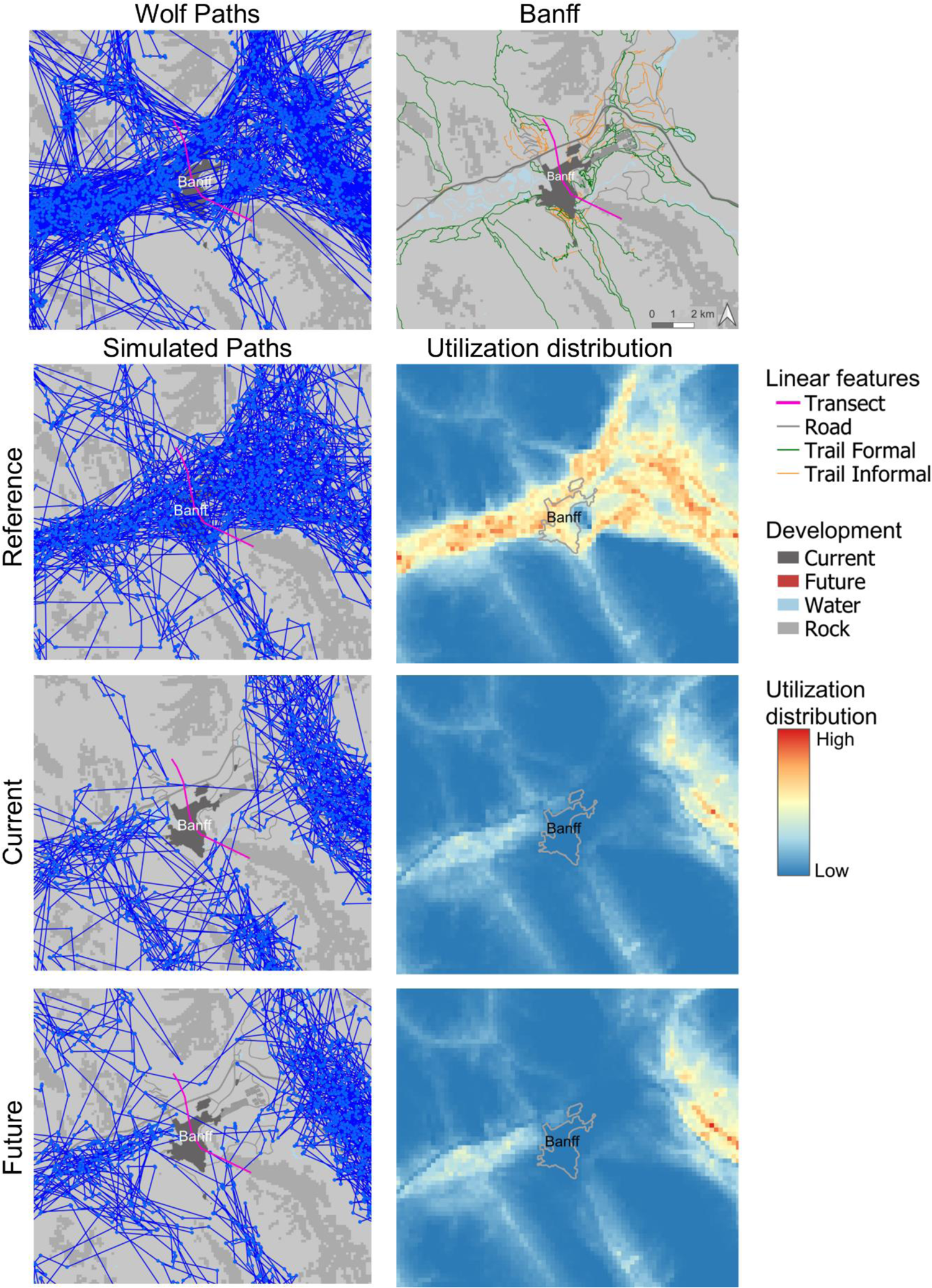
Illustration of our connectivity modeling approach in Figure 1 for one species (wolf) and one season (summer) showing the distribution of observed wolf paths around the town of Banff, a random sample of simulated paths under three land use scenarios, and expected utilization distribution. We used hidden Markov models and SSFs to simulate 200,000 movement paths across a two-month window. We tallied the number of paths that crossed the valley wide transect and calculated connectivity as the ratio of current to reference and future to reference crossing frequencies. We further quantified habitat degradation as changes in the area of high quality habitat relative to reference conditions.

### Statistical analyses

We used a three stage, individual-based modeling approach to quantify carnivore responses to anthropogenic features and connectivity (Figure 1). Here, we provide an overview of our methods and then provide additional details for each step of the analysis. First, we applied hidden Markov models to animal movement data to predict slow versus fast movement states as well as movement parameters and transition probabilities for both movement states. We associated slow states with feeding or resting behaviour, and fast states with travelling behaviour. Second, we integrated movement states into SSFs, such that each SSF contained interactions between movement state, directional persistence, and movement rates. This enabled us to simulate state-specific movements directly from our SSF. We used results of the SSF to assess responses to anthropogenic features. We also developed home-range scale resource selection function (RSF) models to evaluate scale-dependence of SSF models for connectivity evaluation. Third, we used the combination of hidden Markov models and SSFs with covariates to simulate realistic individual-based movements. We simulated movement paths under three landscape conditions reflecting reference, current, and future levels of anthropogenic development. Reference represented a null model of potential habitat with no anthropogenic development. We compared transect crossing rates and UDs from current and future conditions to reference conditions to estimate connectivity and change in the amount of high-quality habitat for each carnivore.

### Movement model

We fit hidden Markov models to grizzly bear and wolf GPS step lengths and turn angles so that we could incorporate movement behaviour into SSFs and to create biologically realistic simulations of animal movement. We used functions from the *moveHMM* package version 1.7 to fit hidden Markov models (Michelot et al. 2016). For each species and season, we fit two-state movement models to reflect slow and fast movements, following previous studies of GPS movement (Fryxell et al. 2008). We used the gamma distribution for step length and the circular von Mises distribution for turn angles (Avgar et al. 2016). We included the cosine of hour as a covariate to allow for diurnal variation in the frequency of slow and fast states. We predicted the probability of being in a fast state for each GPS location, which we then incorporated into the SSF below (Figure 1). We further used parameters from the movement models to simulate movement states, step lengths, and turn angles in the path simulations below.

### Step and resource selection: responses to development across scales

We developed grizzly bear and wolf SSF and RSF to assess how anthropogenic development, topography, and land cover affected seasonal wolf and grizzly bear movement (Figure 1). One of the challenges of interpreting SSFs is that results can depend on sampling scale, i.e., time between locations (Mahoney et al. 2018). To ensure results of our SSF-based movement models were consistent with third-order within home range processes, we developed complementary RSF models for individual wolf and grizzly bears to evaluate potential scale-dependence in SSF results.

The SSF models compared movement and environmental attributes of used steps to matched available locations (Avgar et al. 2016, Signer et al. 2019). We estimated each animal’s movement parameters for step length and turn angle. We sampled from these movement parameters to generate random locations around each used location to sample availability. For each animal’s step (strata *i*) we generated *J* = 10 paired random locations. We extracted covariate vectors ***xij*** for each location and used conditional logistic regression to estimate covariate vector **β**.

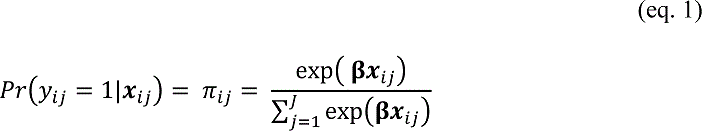

We followed the modelling strategies outlined by Muff et al. (2020) for used-available SSF and RSF designs. We used the Poisson formulation of conditional logistic regression to fit the SSF and included random intercepts for each strata *i*. We accounted for individual animal variability in selection by including random coefficients for explanatory variables. We set weights for used locations to 1.0 and random locations to 1000, and fixed the variance of the random intercept for strata to 10^4^. We used the R packages amt version 0.0.9 (Signer et al. 2019) to define available locations and glmmTMB version 1.0.1 (Brooks et al. 2017) to estimate the models. We visualized the effects of covariates on the relative probability of selection using relative selection strength (RSS) where RSS = exp(**β*x***) (Avgar et al. 2017).

The SSFs included covariates for state-specific speed of travel and directional persistence (Roever et al. 2014, Duchesne et al. 2015). We first predicted the probability of fast state for each used location from the hidden Markov model (section 2.3.1 above, Figure 1). We applied that predicted probability to all paired available locations. Our SSF then included interactions between probability of fast state and the natural logarithm of step length and between probability of fast state and the cosine of turn angle (Avgar et al. 2016). Thus, selection for step lengths (speed) and turn angles (persistence) depended on movement state. The cosine of turn angle reflected a measure of directional persistence with values ranging between −1.0 when animals turned around to 1.0 when they continued in the same direction.

We developed RSF models with a ratio of 1:10 used to available locations and sampled available locations within each individual animal’s 95% minimum convex polygon. We used logistic regression to fit the RSF and included random intercepts for each animal. We accounted for individual animal variability in selection by including random coefficients for explanatory variables. We set weights for used locations to 1.0 and random locations to 1000, and fixed the variance of the random intercept for strata to 10^4^. We used the R packages amt version 0.0.9 (Signer et al. 2019) to define available locations and glmmTMB version 1.0.1 (Brooks et al. 2017) to estimate the models. We used the same explanatory variables for both SSF and RSF models. We visually compared parameter estimates for the SSF and RSF models for consistency in responses to anthropogenic development.

The SSFs and RSFs included environmental and anthropogenic explanatory variables that were previously found to be important predictors of grizzly bear or wolf resource selection in the Canadian Rockies (Nielsen et al. 2006, Hebblewhite and Merrill 2008, Rogala et al. 2011). To minimize collinearity, we removed explanatory variables that had Pearson correlation coefficients > 0.6 and variance inflation factors > 2.0. When two variables were highly correlated, we selected the covariate based on biological relevance and predictive power that we assessed with univariate plots. All models contained the same 17 environmental and anthropogenic covariates. Environmental covariates included five land cover classes, elevation (m), the negative cosine of aspect such that south = 1.0 and north = −1.0, slope (degrees), proximity to forest edge (km), proximity to large patch of vegetated habitat greater than 9 km^2^ (Proctor et al. 2015), and an indicator variable for whether the area had burned since 1960. See Appendix S1: Table S2 for details.

Covariates for anthropogenic development included proximity to towns (km), proximity to campgrounds (km), density of formal trails (km/km^2^, 500 m radius), and indicator variables for whether the animal was on or off trails and the railway. We classified distance to town based on a digitized aerial photograph of buildings and developed areas within towns. We excluded green spaces and golf courses from the town footprint. We predicted that carnivores would select for trails and the railway as travel routes. We predicted that carnivores would avoid areas near towns, campgrounds, and areas of high trail density, especially in summer during peak visitation (Rogala et al. 2011). We lacked direct measures of human activity so we included an interaction between trail density and the natural logarithm of distance to paved road (km). We assumed that trail use would be highest near trail heads along paved roads (Rogala et al. 2011, Zhai et al. 2018). We included an interaction between proximity to town and time of day (cosine of hour) because we expected stronger avoidance of towns during the day compared to the night (Hebblewhite and Merrill 2008). We applied a decay term (1 – exp^-10 * distance^) to the distance covariates so that the influence of these features had an asymptote near 500 m (Shepherd and Whittington 2006, Rogala et al. 2011). We scaled all other continuous covariates by their mean and standard deviation to improve model convergence.

Animal resource selection and responses to anthropogenic development can vary seasonally. Thus, we defined four seasons and created separate SSF models for each species and season. We defined seasons based on animal movement, plant phenology, and human visitation rates to BNP. We classified *Spring* as May and June which included plant emergence, ungulate parturition, grizzly bear mating, wolf denning, and moderate levels of human activity; *Summer* as July and August during the height of berry season and peak visitation; and *Fall* as September and October when plants have senesced and the study area received moderate levels of visitation, and *Winter* as November through April for wolves with lower levels of visitation in backcountry areas and high levels of visitation near ski hills and towns.

### Connectivity and habitat degradation

We simulated individual-based carnivore movements from our SSFs across three landscape scenarios, from which we estimated connectivity and changes in the amount of high quality habitat (Figure 1). We used a combination of the hidden Markov models and SSFs to simulate carnivore movements throughout the study area (Figure 1, Appendix S1: Figure 1). We simulated 200,000 paths within the 17,000 km^2^ study are for each species, season, and landscape scenario. We selected random start locations and initial directions of travel. For each path, we sampled *s* = 720 movement states (steps, *s*) with two-hour fix interval across *t* = 60 days from the hidden Markov models. We chose 60 days to match the duration of spring, summer, and fall seasons used in the step selection analyses. For each step, we sampled *n* = 20 step lengths and turn angles from the state-specific movement parameters. We extracted environmental attributes of the proposed locations and used the combination of environmental attributes and movement parameters to calculate probability of use conditional on the 20 sample locations (equation 1). We probabilistically selected one of the proposed locations and continued to the next step. We repeated this process for all steps in the path.

The study occurred in a rugged environment where steep, rocky mountain ranges can influence animal movements. We therefore defined unavailable habitat as barren landscapes with slopes ≥ 35 degrees, which were used by grizzly bears and wolves 1.9 and 0.2% of the time, respectively. We also classified towns and developed areas as unavailable habitat. To create realistic movement paths, we reduced the probability of simulated steps jumping across mountain ranges and towns by sampling four equidistant locations along proposed steps. We rejected steps if any of those locations occurred in the unavailable habitat. We minimized boundary effects on spatial predictions of use by terminating paths when > 40% of the proposed steps occurred outside the study area and by setting the study area boundary > 30 km from the towns of Banff and Canmore. Finally, start locations could occur in poor quality habitat, so we removed the first twelve (*t* = 1 day) steps from each path while paths oriented to higher quality habitat.

We simulated animal movements for three scenarios with varying levels of anthropogenic development: reference, current, and future. First, we removed the effect of towns, roads, and the railway from SSFs when simulating paths under reference conditions, which we used as a null model of movement (Heinemeyer et al. 2019, Brennan et al. 2020). In reality, First Nations have occupied the study area for over 11,000 years (Langemann 2011) and the reference condition underestimated the historical effects of human activity. Second, we simulated animal movements under current conditions from which we developed our SSFs. Finally, we simulated animal movements under one future scenario with expanded development and trail density. We modified the town of Canmore’s developed footprint to reflect residential and business development proposals in the 2020 Smith Creek and Three Sisters area structure plans (QuantumPlace Developments Ltd. 2020b, a). We excluded green spaces and golf courses from the developed footprint given that carnivores can use these areas for movement. The developed footprint for the town of Banff is legally fixed under the National Parks Act and is not expected to increase. However, like many mountain towns, the creation and intensity of use on informal trails has increased near Banff and Canmore over the last ten years. We, thus added an inventory of informal trails to the existing formal trail network and updated metrics of trail density. Increased use of existing and new recreational trails has the potential to reduce wildlife connectivity (Gutzwiller et al. 2017). We simulated animal movements with the updated town and trail layers to estimate future connectivity.

To calculate connectivity, we created digital, cross-valley transects through the towns of Banff and Canmore (Figure 2, Figure 3). We aligned transects so that they crossed the narrowest movement corridors under current condition, where the combination of rugged topography and development created pinch points to movement. We counted both the number of simulated paths and individual steps that crossed transects on the north and south sides of the valley. We used number of unique paths that crossed the transects as our metric of connectivity to reflect the population level value of corridors. We calculated connectivity as 100 * *n_cross_* / *n_reference_*, where *n_cross_* was the number of unique paths that crossed in current or future conditions and *n_reference_* was the number of unique paths that crossed under reference conditions with no anthropogenic development. We evaluated how connectivity changed with species, seasons, and time period.

**Figure 3.**
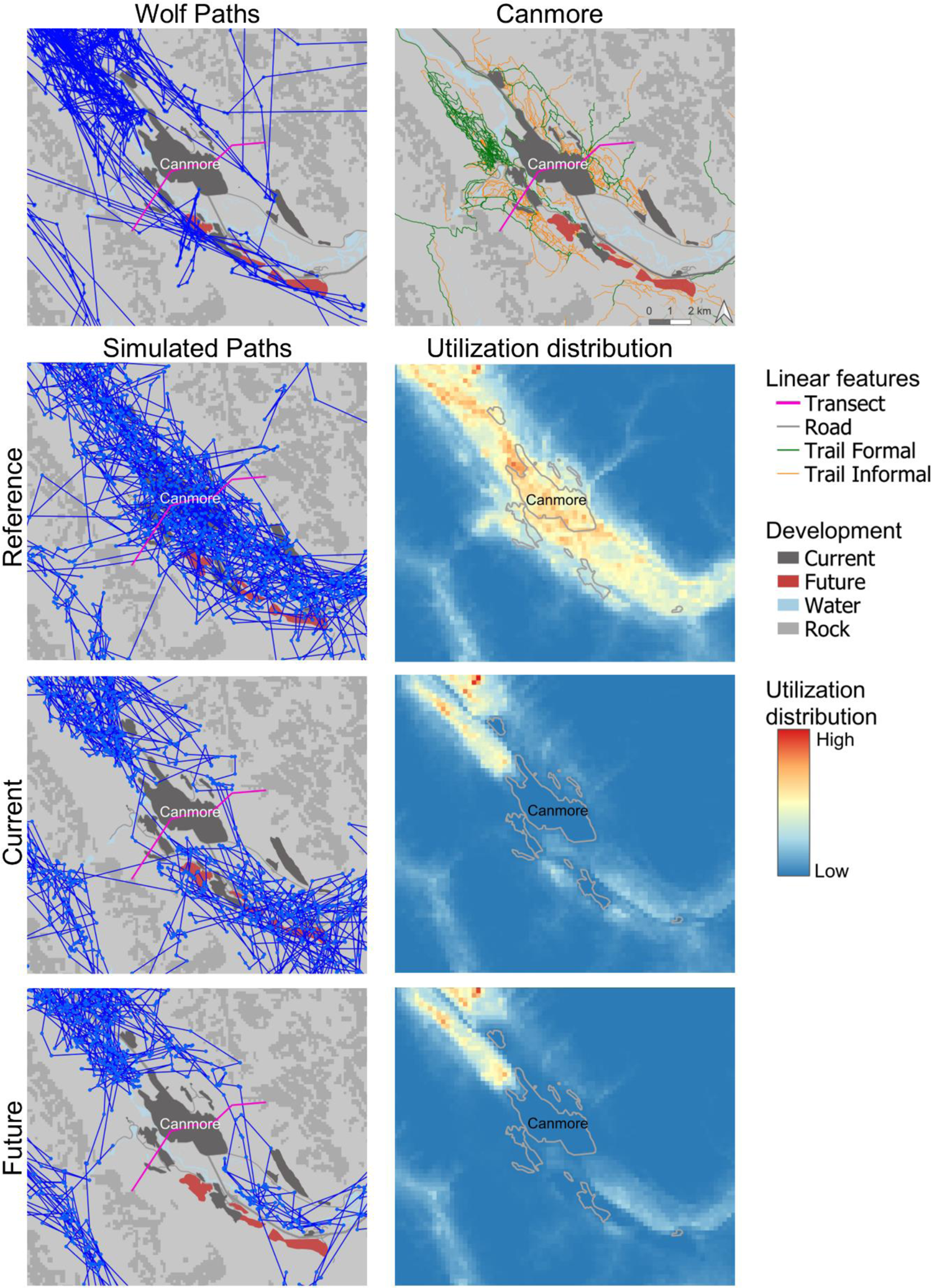
Illustration of our connectivity modeling approach in Figure 1 for one species (wolf) and one season (summer) showing the distribution of observed wolf paths around the town of Canmore, a random sample of simulated paths under three land use scenarios, and expected utilization distribution.

Finally, we examined the effects of anthropogenic development on the amount of high quality habitat available to carnivores. We calculated UDs as the number of simulated locations that occurred within each 210 x 210 m^2^ grid cell and then divided the tallies by the number of total simulated locations (Signer et al. 2017). We classified reference UDs into three equal area bins representing low, medium, and high quality habitat. We applied the same break points and habitat classifications to UDs from the current and future scenarios. We then calculated changes in the amount of high quality habitat. We focussed our analysis within a five km radius of the Trans Canada Highway between Banff and Canmore (366 km^2^). The five km radius represented the 0.99 and 0.95 quantiles of grizzly bear and wolf step lengths, respectively, and the focal study approximately covered the peak to peak width of the Bow Valley. We calculated the proportion of high quality habitat degraded due to anthropogenic development relative to reference conditions (Heinemeyer et al. 2019). For example, our calculation of habitat degradation under current conditions was (*AreaHigh_Reference_* – *AreaHigh_Current_*)/*TotalArea*, whereby *AreaHigh* represented the area of high quality habitat and *TotalArea*, represented the total area of the focal study. Our metric of habitat degradation thus accounted for both decreased UDs near anthropogenic developments and concurrent increased UDs as simulated animals spent more time in less developed portions of the landscape. We visually evaluated how habitat degradation varied with species, seasons, and time period.

## Results

### Movement state

We analysed GPS data from 34 grizzly bears (19 females, 15 males, 72,217 locations) and 33 wolves (13 females, 20 males, 84,434 locations; Appendix S1: Figure S1 – S3). Hidden Markov models revealed that grizzly bears and wolves spent a similar proportion of time in their fast state (*p* = 0.64 and 0.60 respectively). Grizzly bears and wolves had the same median step lengths for slow steps (16 m) (Appendix S2: Table S1). Wolf fast steps (median = 1270 m) were on average 2.5 times longer than grizzly bear fast steps (median = 496 m). Grizzly bears had a much stronger diurnal cycle of movement states than wolves (Figure 4). Grizzly bears increased their proportion of time in slow states at night. Wolves had a weaker and sometimes opposite diurnal cycle. Wolves increased the proportion of time in slow states at night during fall and winter only. Wolves increased the proportion of time in fast states at night during spring and summer, which coincided with the longest days of the year.

**Figure 4.**
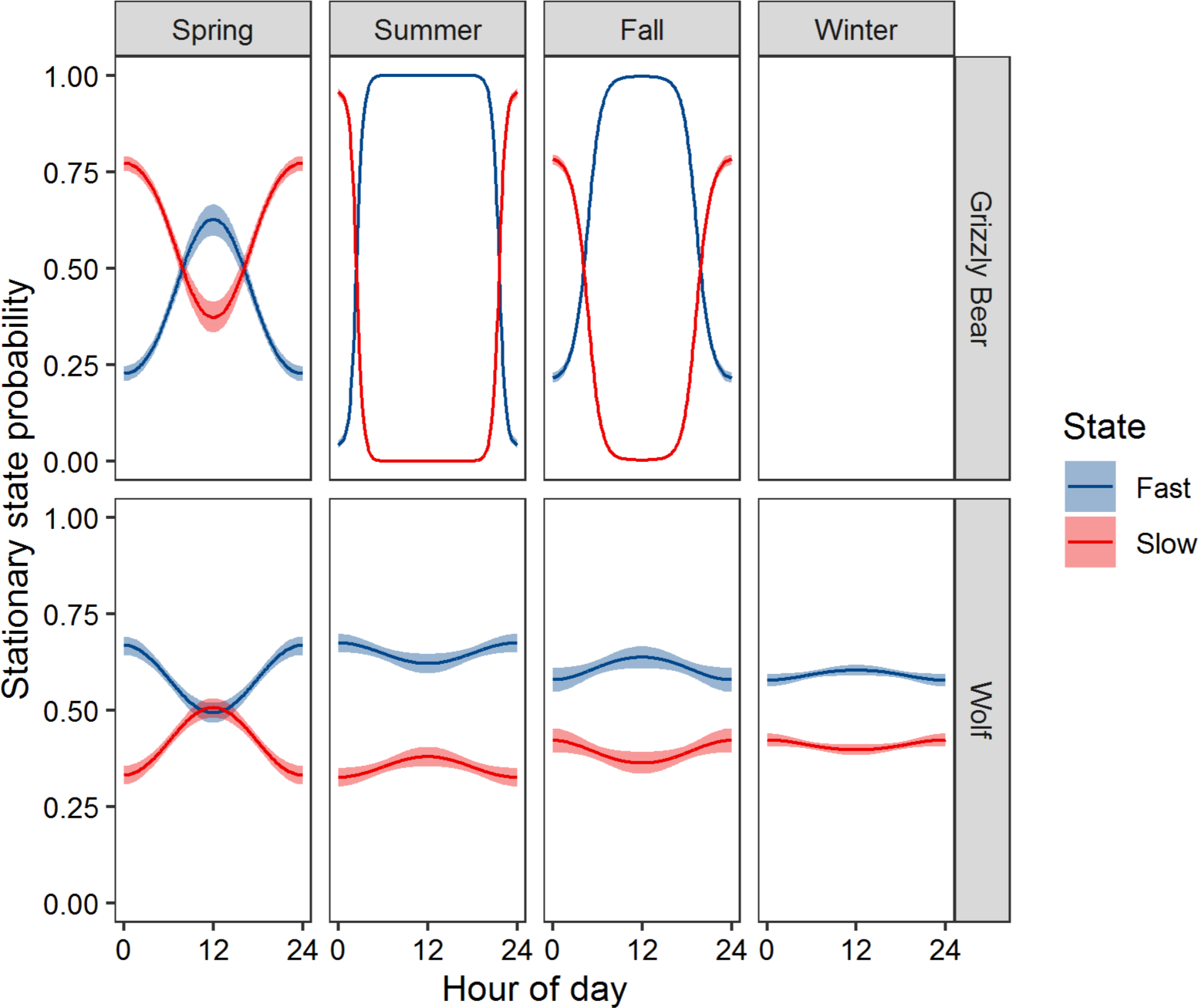
Grizzly bear and wolf movement state probabilities (slow versus fast movements) and 95% CI’s depended on time of day. Movement states were predicted from hidden Markov models developed from GPS locations. Slow and fast states are interpreted to represent feeding-resting and travel behaviours, respectively.

### Step and resource selection: responses to development across scales

As expected, we found that wolves and grizzly bears generally avoided areas with high levels of human activity in all seasons (Figure 5, Figure 6, Supplementary Table S2). Both species strongly avoided areas near towns (median β = 1.11, range from 0.44 to 2.08) and 95% CI’s excluded zero on 6 of the 7 models (Figure 4). Grizzly bear and wolf responses to areas near town changed slightly at night, though the effect size was small compared to avoidance of towns in general. Parameter estimates for distance to town were >10 times larger than parameter estimates for the distance to town by night time interaction (Figure 5, Figure 6). Grizzly bear avoidance of towns diminished at night in all seasons (e.g., summer β = −0.05, SE = 0.01). Wolf response to towns diminished at night during winter (β = −0.05, SE = 0.01), but strengthened during summer (β = −0.05, SE = 0.01), and fall (β = −0.05, SE = 0.01). Grizzly bears and wolves avoided areas with high trail density near paved roads (median β = −0.93, range from −1.92 to 0.151) with all but one estimate being less than zero and five out of seven models with 95% CI’s that excluded zero. Carnivore responses to trail density tapered with distance to paved roads (Figure 6). Wolves avoided areas near vehicle accessible campgrounds during summer when campgrounds were most active (e.g., summer β = 1.00, SE = 0.41). Grizzly bears avoided areas near campgrounds during the fall but not during the summer berry season nor in the spring.

**Figure 5.**
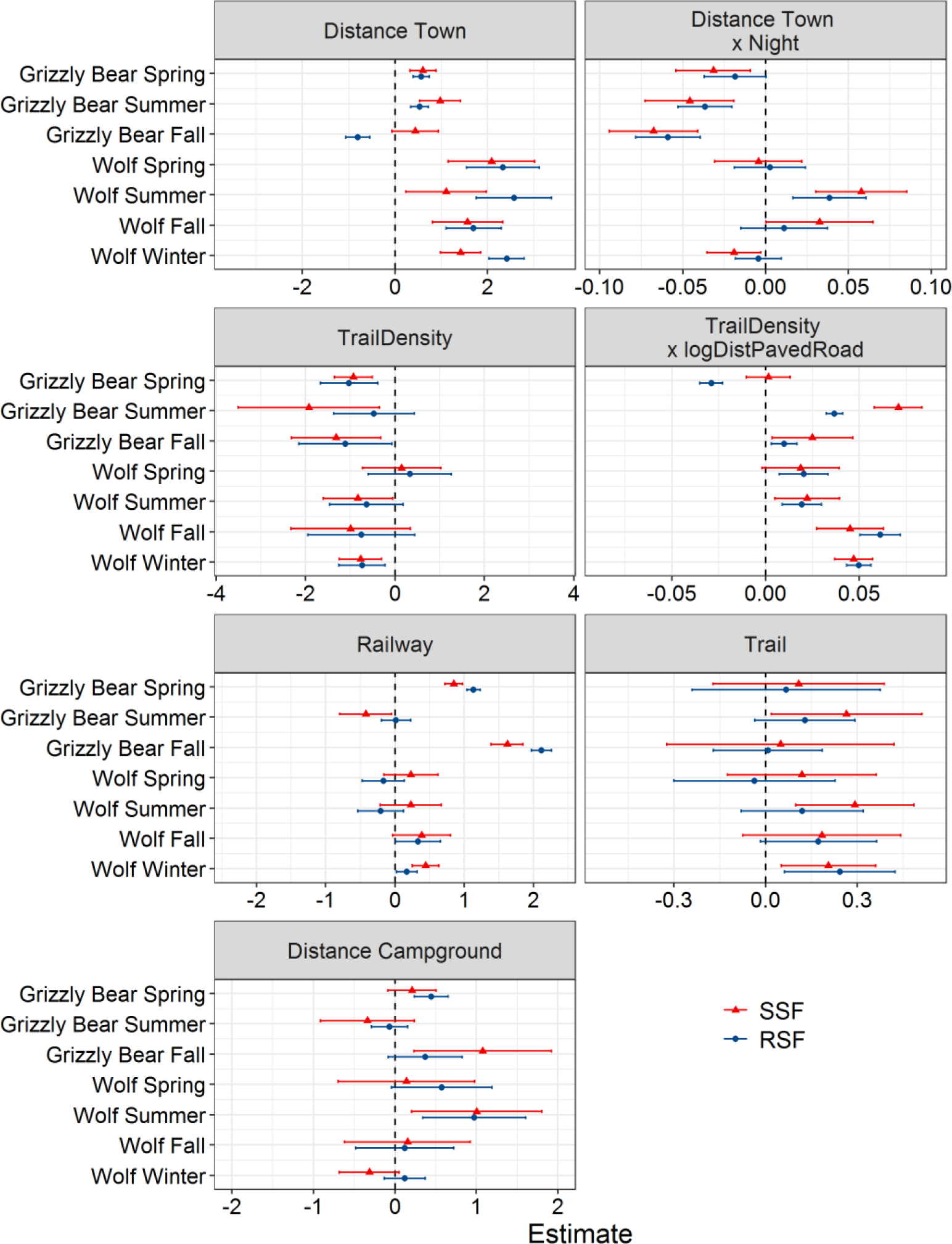
Human use related parameter estimates and 95% CI’s from grizzly bear and wolf SSF and RSF models. We created separate models for each species and season. Positive values reflect selection for high values of the covariate.

**Figure 6.**
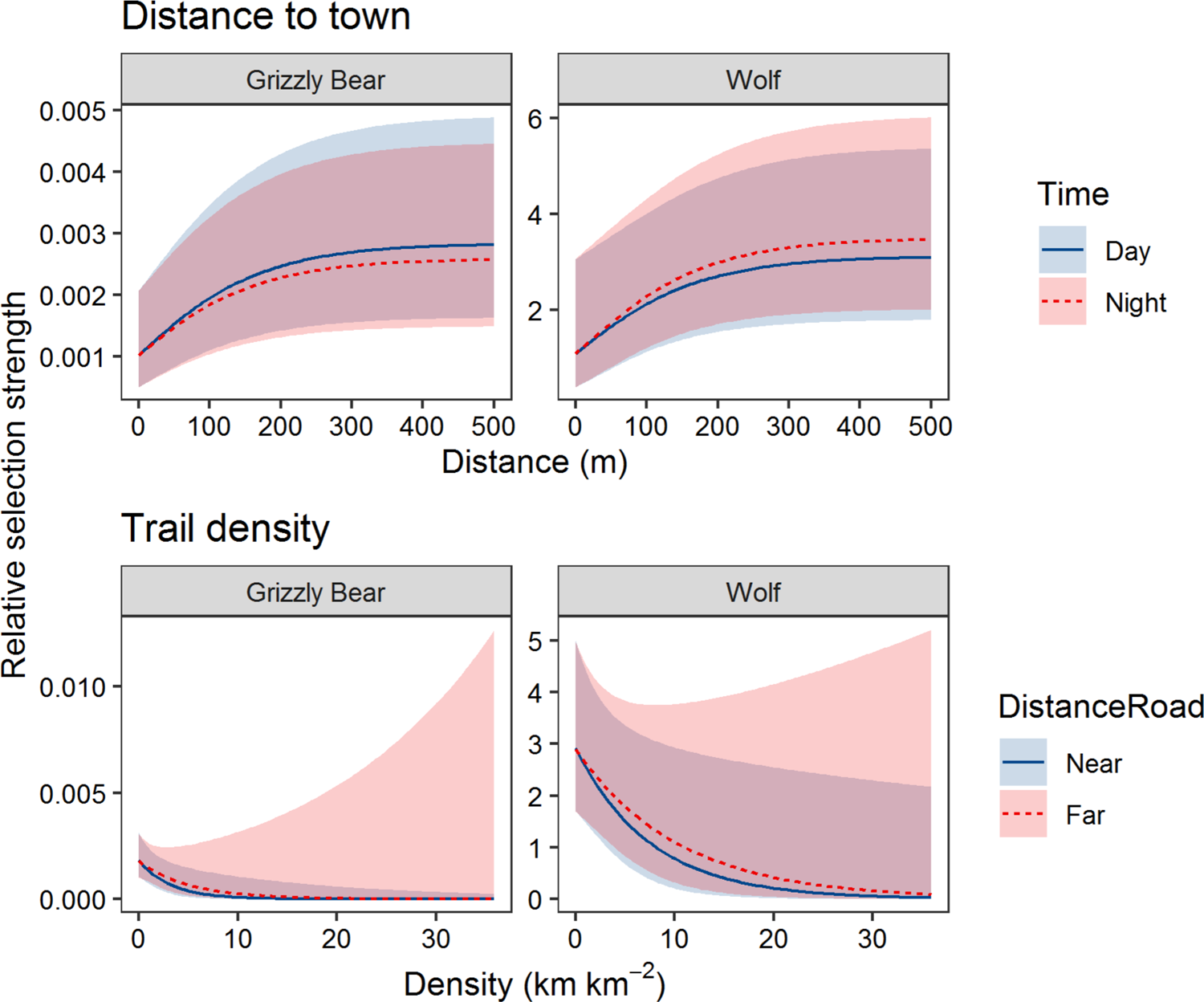
Grizzly bear and wolf relative selection strength and 95% CI’s as a function of distance to town and trail density during summer. We calculated relative selection strength by creating predictions from SSFs while holding all variables constant at their mean except for distance to town, trail density, time of day (Day = 1200 hours, Night = 2400 hours), and distance to paved road (Near = 0 km and Far = 20 km).

Grizzly bears and wolves weakly selected trails during all seasons with the strongest selection for grizzly bears in summer and for wolves in winter. Grizzly bears selected the railway during the spring and fall and avoided the railway during the summer berry season. Wolves strongly selected the railway during winter and weakly selected the railway at other times of the year.

RSF models had similar parameter estimates compared to SSF models confirming minimal scale-dependence of our SSF results (Figure 5, Appendix S2: Table S3, Table S4). Most (84%) of the SSF and RSF anthropogenic parameters had the same positive or negative sign.

Most sign differences occurred for parameters with 95% CI’s that overlapped zero. From a management perspective, the biggest difference in parameter estimates was that the grizzly bear fall SSF suggested weak avoidance of areas near town (β = 0.44, SE = 0.26), whereas the RSF suggested grizzly bears selected areas near town (β = −0.81, SE = 0.13). Otherwise, all other parameter estimates for distance to town were positive. Overall, the RSF results supported the scale-independence of our SSF results regarding carnivore avoidance of areas near towns and areas with high trail density.

### Connectivity and habitat degradation

Simulated paths under reference, current, and future land use scenarios had similar movement attributes compared observed paths (e.g., Figure 2, Figure 3). Both simulated and observed paths contained series of short steps with high turn angles interspersed with long distance movements with strong directional persistence. Under reference conditions, simulated paths were concentrated in the valley bottoms and used areas within the current footprint of towns. The combination of towns and rugged topography constrained the movements of both observed and simulated paths under current and future scenarios. This resulted in UDs with low frequencies of occurrence near towns and areas of high trail density and high UDs in more remote areas of the Bow Valley (Appendix S1: Figures S4 - S5).

Grizzly bear and wolf connectivity across digital transects on the north and south sides of Banff and Canmore ranged between 7 and 45% under current conditions with mean values of 21% for grizzly bears and 25% for wolves (Figure 7). Grizzly bear and wolf connectivity further decreased from current to future conditions an average of 6% and 5% respectively (range = 0 to 13%). Connectivity for grizzly bears and wolves was highest in the spring. Grizzly bear connectivity was lowest in the summer, whereas wolf connectivity was lowest in the fall and winter. Grizzly bear connectivity was on average higher along the northern transects compared to the southern transects. Wolf connectivity was highest on the northern side of Banff and lowest on the southern side of Canmore.

**Figure 7.**
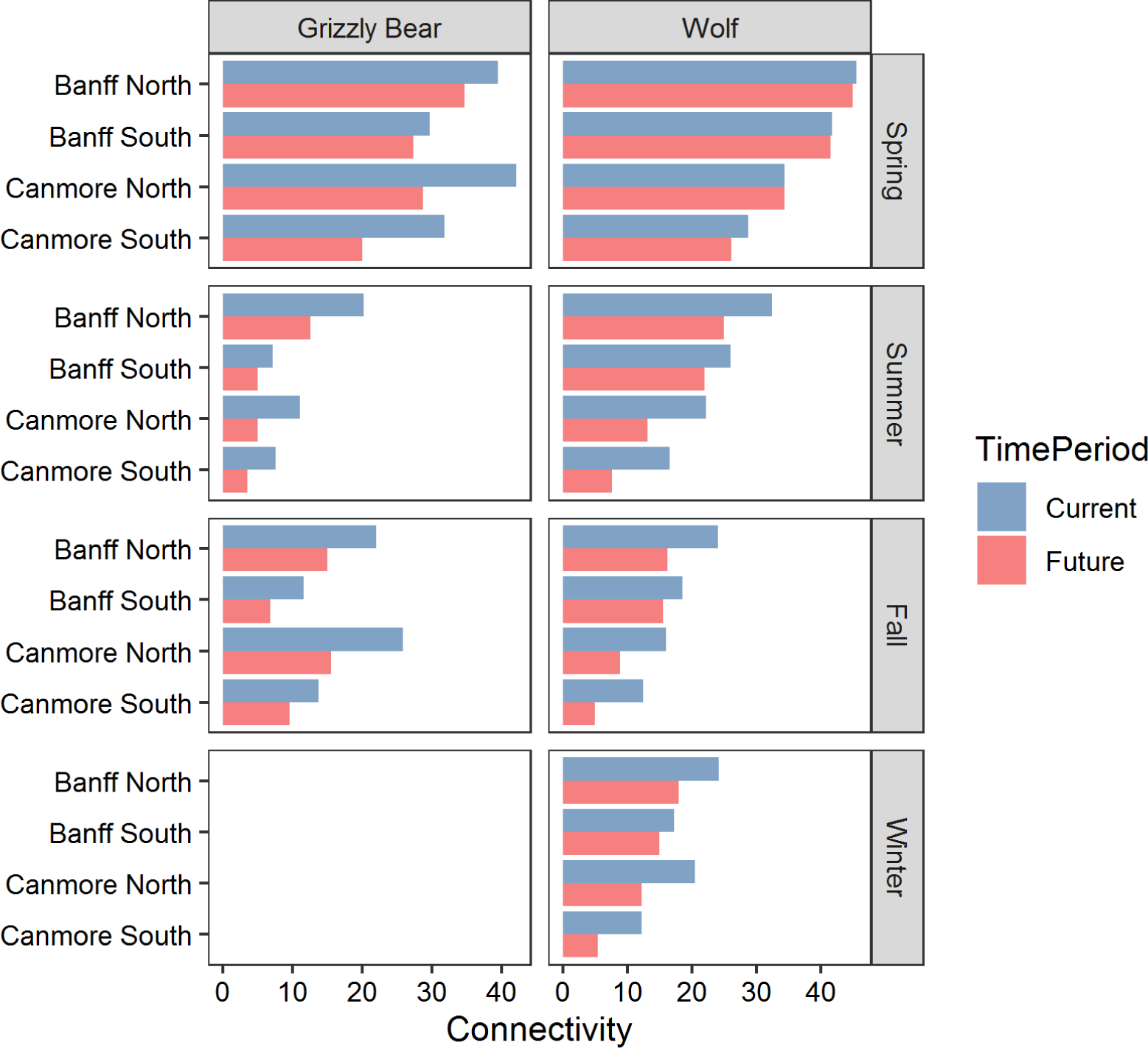
Connectivity estimates for grizzly bears and wolves around the towns of Banff and Canmore under current and future footprints of anthropogenic development. We estimated connectivity by comparing the number of simulated paths that crossed transects under current and future conditions to crossing rates from reference conditions. We simulated 200,000 paths for each species, season, and time period. On average, connectivity decreased from Current to Future for grizzly bears by 6.5%, and, wolves by 5.1%. Grizzly bears have no connectivity estimates while they hibernation in winter.

Grizzly bears and wolves UDs showed high intensity of use through the valley bottoms including areas near Banff and Canmore under reference conditions (Figure 2, Figure 3, Supplementary Figures S4 and S5). UDs under current and future conditions showed a cumulative decrease in use in and around the towns. The decrease in use near towns was offset by increased use in more remote areas of the valley. The proportion of high quality habitat degraded due to anthropogenic development increased from current (mean = 0.145, range = 0.088 to 0.183) to future conditions (mean = 0.164, range = 0.126 to 0.198; Figure 8). Habitat degradation was highest in summer and lowest in the spring and fall for grizzly bears. Habitat degradation was high for wolves in the summer, fall, and winter, and lowest in the spring.

**Figure 8.**
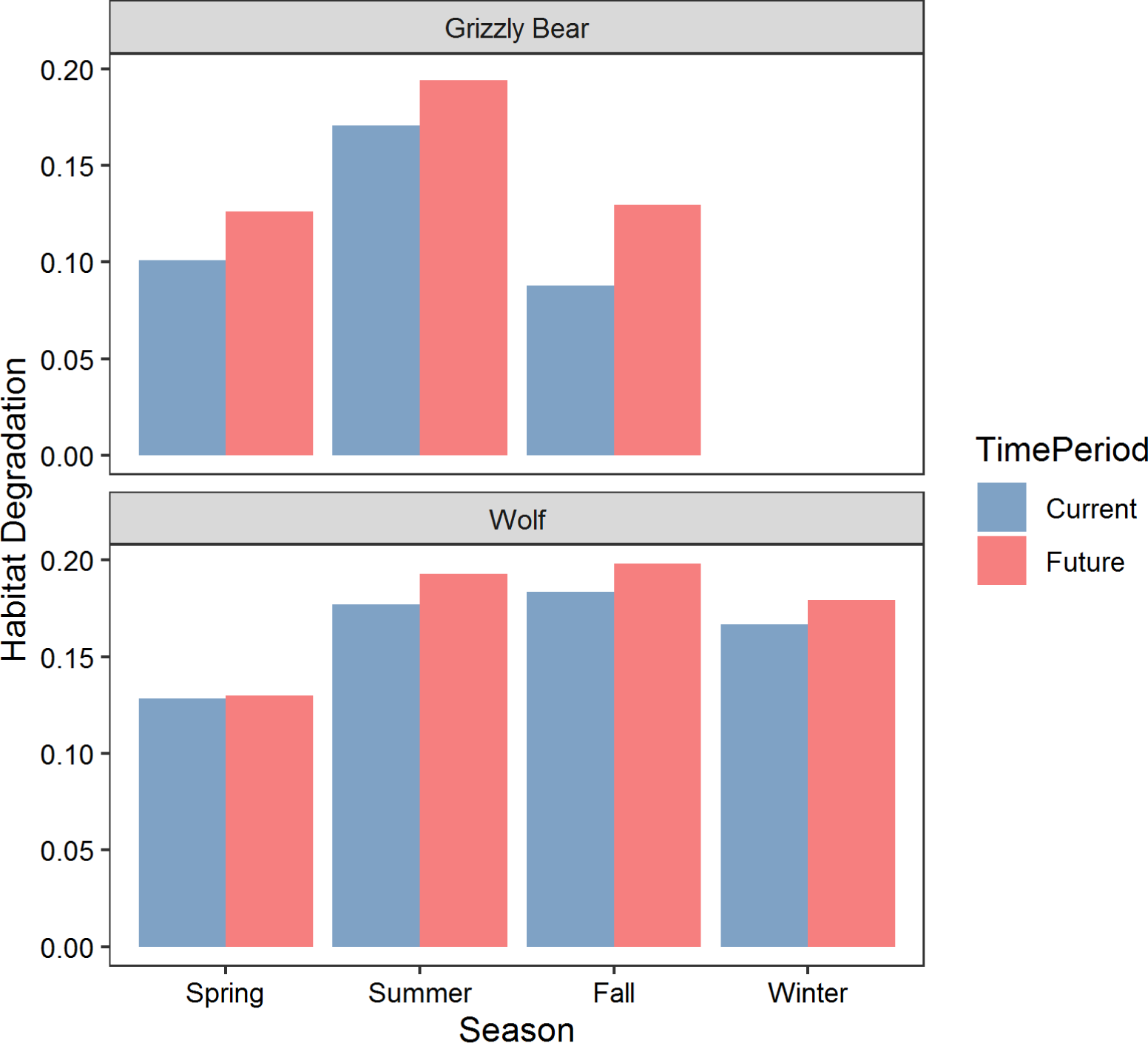
Habitat degradation measured as the decrease in proportion of high quality habitat from reference to current and future time periods. We estimated habitat degradation for the Bow Valley between Banff and Canmore, Alberta, Canada

## Discussion

Our study used a combination of hidden Markov models and SSFs to estimate utilizations, connectivity, and changes in the amount of high quality due to current and future estimates of anthropogenic development. Our approach assessed the cumulative effects of multiple anthropogenic features on carnivore movements and connectivity around the towns of Banff and Canmore, Alberta, which averaged 23% relative to reference conditions. A scenario of future development and trail expansion would further reduce connectivity by an average of 5 to 6%. By using empirically-derived parameters in an individual-based simulation, our approach offers a tangible response variable for scientists to convey to land use decision makers. For example, rather than translating how ‘current density’ may change under different scenarios, we are able to estimate a percent change in the number of animals moving through a corridor under predicted land use scenarios.

The main advantages of our approach are as follows: 1) simulated movements directly from an SSF with multiple behavioural states helped create realistic movement paths where movement decisions were based on resource selection parameters; 2) our approach avoided transforming SSFs into resistance surfaces used for circuit-theory and cost-distance analyses, which have an weak theoretical link to movement ecology; and 3) estimates for changes in UDs and connectivity have a simpler interpretation and a tighter link to movement probabilities compared to least-cost and circuit theory based estimates of connectivity. We chose to estimate connectivity by comparing transect crossing rates of simulated paths through narrow movement corridors, which in our study area are of conservation concern. Our approach could easily be expanded to examine the frequency with which paths travel between habitat patches (e.g. Suraci et al. 2020), between summer and winter ranges (Merkle et al. 2019), across highways with increased mortality risk (e.g. Quaglietta et al. 2019), or through other areas of conservation concern.

Our study supports the growing body of research showing that wildlife avoid some forms of human activity (e.g. Gaynor et al. 2018, Tucker et al. 2018, Nickel et al. 2020), which can lead to habitat fragmentation and reductions in connectivity (e.g. Bischof et al. 2017, Hilty et al. 2020, Suraci et al. 2020). Given the global growth in human activity adjacent to protected areas (Wittemyer et al. 2008), and concurrent impacts of growing recreation in these landscapes (Gutzwiller et al. 2017), our approach and results emphasize the importance of cumulative effects assessment in regions surrounding parks and protected areas.

Numerous studies have found that grizzly bears (Chetkiewicz and Boyce 2009, Morales-González et al. 2020) and wolves (Hebblewhite and Merrill 2008, Rogala et al. 2011, Anton et al. 2020) avoid human activity, which can contribute to the fragmentation of populations (Proctor et al. 2012, Bischof et al. 2017). However, few studies have compared the behaviour of the two species. Wolves in our study exhibited stronger avoidance of towns, similar responses to trails, and weaker selection for the railway compared to grizzly bears. Grizzly bears and wolves had higher connectivity estimates in spring, which coincided with lower levels of human activity and a concentration of food resources and wolf movements to and from den sites in valley bottoms. Interestingly, wolf connectivity estimates were slightly higher than grizzly bear connectivity estimates. One reason for this disconnect could be that wolves had faster speeds of travel, fewer steps were required to traverse corridors, and perhaps simulated steps could more easily jump across towns. Parameterizing models using path selection functions or collecting finer resolution GPS data could help reduce the probability of paths crossing inhospitable features. For instance, path selection functions can sometimes produced stronger regression coefficients and better connectivity models compared to SSFs (Zeller et al. 2015, Zeller et al. 2018). Further, we did not assess how individual variability in animal responses to anthropogenic development affected connectivity (Muff et al. 2020). Simulating movements from random coefficients could highlight estimates of connectivity for both wary and habituated animals and could help identify areas likely to have high levels of human wildlife conflict (Buchholtz et al. 2020, Lamb et al. 2020).

Two limitations of our study bear further consideration for similar work in the future. First, we lacked direct measurements of human activity on trail networks (Alberta Environment and Parks 2018). Because carnivores typically avoid encounters with people rather than linear features, our lack of high resolution trail use data likely reduced the effect size and precision of parameter estimates for trail density. Estimates of recreational activity could be improved by directly tracking individual movements (Heinemeyer et al. 2019), inferring activity from mobile device data (Corradini et al. 2021), or modelling spatial and temporal trends in trail use (Ladle et al. 2019). Better estimates of recreational activity would improve our understanding of how recreational activity affects wildlife movement and our ability to manage human-wildlife coexistence (Rogala et al. 2011, Naidoo and Burton 2020). Second, our data consisted of animal movements within established home ranges rather than dispersal or nomadic movements that are important for landscape-scale connectivity (Fattebert et al. 2015). Other studies suggest animals select different resources and may have increased tolerance for human activity when dispersing when compared to movement within the home range. For instance, resistance models for Iberian lynx under-estimated connectivity when they were developed using GPS data from within home range movements (Blazquez-Cabrera et al. 2016). Further development and evaluation of connectivity models using dispersal data would be important when evaluating connectivity between isolated populations (Zeller et al. 2018).

Restoration actions, such as removal of human footprint, managing or consolidating recreational activity, and trail closures have potential to improve habitat quality and connectivity. Wildlife increased their use of corridors and degraded habitat following reductions in human activity, both in our ecosystem (Duke et al. 2001, Shepherd and Whittington 2006, Whittington et al. 2019) and around the world (Ngoprasert et al. 2017). For example, early work in our study area demonstrated positive wildlife connectivity consequences of removing recreational footprint in the Cascade wildlife corridor on the north side of the Banff town site (Duke et al. 2001), and positive effects of a temporal road closure on wildlife habitat quality (Whittington et al. 2019).

Our approach for simulating animal movements and assessing connectivity could be applied to assess the effects of potential restoration actions on fine-scale connectivity (Wang et al. 2014, Mariela et al. 2020, Suraci et al. 2020). Simulations and restoration actions could focus on highway mitigations (Quaglietta et al. 2019), reductions in trail density, permanent closures, seasonal closures, or temporal closures (Whittington et al. 2019). In the face of global increases in human activity, especially surrounding parks and protected areas (Wittemyer et al. 2008), proactive habitat protection and restoration actions will be required to maintain habitat quality and connectivity for wide ranging wildlife (Hilty et al. 2020).

## Acknowledgements

Parks Canada, the University of Montana, the University of Alberta, and Canadian Pacific Railway contributed to the collection of grizzly bear and wolf GPS data. MH acknowledges funding from NSF LTREB grant # 1556248. We thank J. Merkle for provided advice on step selection analyses and B. Hunt, A. Forshner, and S. Williams for their insights on earlier versions of the manuscript. Parks Canada provided financial support for data collection and analysis.

## Supporting Information

Appendix S1. Maps of study area, observed GPS locations, and predicted utilization distributions from step selection function models.

Appendix S2: Tables of parameter estimates from movement models, step selection functions, and resource selection functions.

## Data availability

We will submit GPS movement data, scripts to fit hidden Markov models, step selection functions, and simulations under current conditions to Data Dryad.

## Literature cited

1. Abrahms, B., S. C. Sawyer, N. R. Jordan, J. W. McNutt, A. M. Wilson, and J. S. Brashares. 2017. Does wildlife resource selection accurately inform corridor conservation? Journal of Applied Ecology 54:412–422.

2. Alberta Environment and Parks. 2018. Human-wildlife coexistence: recommendations for improving human-wildlife coexistence in the Bow Valley.

3. Anton, C. B., D. W. Smith, J. P. Suraci, D. R. Stahler, T. P. Duane, and C. C. Wilmers. 2020. Gray wolf habitat use in response to visitor activity along roadways in Yellowstone National Park. Ecosphere 11:e03164.

4. Avgar, T., S. R. Lele, J. L. Keim, and M. S. Boyce. 2017. Relative selection strength: quantifying effect size in habitat- and step-selection inference. Ecology and Evolution 7:5322–5330.

5. Avgar, T., J. R. Potts, M. A. Lewis, and M. S. Boyce. 2016. Integrated step selection analysis: bridging the gap between resource selection and animal movement. Methods in Ecology and Evolution 7:619–630.

6. Benz, R. A., M. S. Boyce, H. Thurfjell, D. G. Paton, M. Musiani, C. F. Dormann, and S. Ciuti. 2016. Dispersal ecology informs design of large-scale wildlife corridors. PLoS ONE 11:e0162989.

7. Bischof, R., S. M. J. G. Steyaert, and J. Kindberg. 2017. Caught in the mesh: roads and their network-scale impediment to animal movement. Ecography 40:1369–1380.

8. Blazquez-Cabrera, S., A. Gastón, P. Beier, G. Garrote, M. Á. Simón, and S. Saura. 2016. Influence of separating home range and dispersal movements on characterizing corridors and effective distances. Landscape Ecology 31:2355–2366.

9. Brennan, A., P. Beytell, O. Aschenborn, P. Du Preez, P. J. Funston, L. Hanssen, J. W. Kilian, G. Stuart-Hill, R. D. Taylor, and R. Naidoo. 2020. Characterizing multispecies connectivity across a transfrontier conservation landscape. Journal of Applied Ecology 57:1700–1710.

10. Brooks, M. E., K. Kristensen, K. J. van Benthem, A. Magnusson, C. W. Berg, A. Nielsen, H. J. Skaug, M. Machler, and B. M. Bolker. 2017. glmmTMB balances speed and flexibility among packages for zero-inflated generalized linear mixed modeling. The R journal 9:378–400.

11. Buchholtz, E. K., A. Stronza, A. Songhurst, G. McCulloch, and L. A. Fitzgerald. 2020. Using landscape connectivity to predict human-wildlife conflict. Biological Conservation 248:108677.

12. Buderman, F. E., M. B. Hooten, J. S. Ivan, and T. M. Shenk. 2018. Large-scale movement behavior in a reintroduced predator population. Ecography 41:126–139.

13. Calabrese, J. M., and W. F. Fagan. 2004. A comparison-shopper’s guide to connectivity metrics. Frontiers in Ecology and the Environment 2:529–536.

14. Chetkiewicz, C. L. B., and M. S. Boyce. 2009. Use of resource selection functions to identify conservation corridors. Journal of Applied Ecology 46:1036–1047.

15. Corradini, A., M. Randles, L. Pedrotti, E. van Loon, G. Passoni, V. Oberosler, F. Rovero, C. Tattoni, M. Ciolli, and F. Cagnacci. 2021. Effects of cumulated outdoor activity on wildlife habitat use. Biological Conservation 253:108818.

16. Duchesne, T., D. Fortin, and L.-P. Rivest. 2015. Equivalence between step selection functions and biased correlated random walks for statistical inference on animal movement. PLoS ONE 10:e0122947.

17. Duke, D. L., M. Hebblewhite, P. C. Paquet, C. Callaghan, and M. Percy. 2001. Restoring a large carnivore corridor in Banff National Park. Pages 261-275 in D. Maehr, R. F. Noss, and J. Larkin, editors. Large mammal restoration: ecological and social challenges in the 21st century. Island Press, Washington, DC.

18. Fattebert, J., H. S. Robinson, G. Balme, R. Slotow, and L. Hunter. 2015. Structural habitat predicts functional dispersal habitat of a large carnivore: how leopards change spots. Ecological Applications 25:1911–1921.

19. Ford, A. T., E. J. Sunter, C. Fauvelle, J. L. Bradshaw, B. Ford, J. Hutchen, N. Phillipow, and K. J. Teichman. 2020. Effective corridor width: linking the spatial ecology of wildlife with land use policy. European Journal of Wildlife Research 66:69.

20. Fortin, D., H. L. Beyer, M. S. Boyce, D. W. Smith, T. Duchesne, and J. S. Mao. 2005. Wolves influence elk movements: behavior shapes a trophic cascade in Yellowstone National Park. Ecology 86:1320–1330.

21. Fryxell, J. M., M. Hazell, L. Börger, B. D. Dalziel, D. T. Haydon, J. M. Morales, T. McIntosh, and R. C. Rosatte. 2008. Multiple movement modes by large herbivores at multiple spatiotemporal scales. Proceedings of the National Academy of Sciences 105:19114–19119.

22. Fullman, T. J., R. R. Wilson, K. Joly, D. D. Gustine, P. Leonard, and W. M. Loya. 2021. Mapping potential effects of proposed roads on migratory connectivity for a highly mobile herbivore using circuit theory. Ecological Applications 31:e2207.

23. Gaynor, K. M., C. E. Hojnowski, N. H. Carter, and J. S. Brashares. 2018. The influence of human disturbance on wildlife nocturnality. Science 360:1232–1235.

24. Gutzwiller, K. J., A. L. D’Antonio, and C. A. Monz. 2017. Wildland recreation disturbance: broad-scale spatial analysis and management. Frontiers in Ecology and the Environment 15:517–524.

25. Hebblewhite, M. 2005. Predation by wolves interacts with the North Pacific Oscillation (NPO) on a western North American elk population. Journal of Animal Ecology 74:226–233.

26. Hebblewhite, M., and E. Merrill. 2008. Modelling wildlife–human relationships for social species with mixed-effects resource selection models. Journal of Applied Ecology 45:834–844.

27. Hebblewhite, M., and E. H. Merrill. 2011. Demographic balancing of migrant and resident elk in a parially migratory population through forage-predation tradeoffs. Oikos 120:1860–1870.

28. Hebblewhite, M., M. Percy, and E. H. Merrill. 2007. Are all global positioning system collars created equal? Correcting habitat-induced bias using three brands in the Central Canadian Rockies. Journal of Wildlife Management 71:2026–2033.

29. Hebblewhite, M., C. A. White, C. Nietvelt, J. M. McKenzie, T. E. Hurd, J. M. Fryxell, J. M. Bayley, and P. C. Paquet. 2005. Human activity mediates a trophic cascade caused by wolves. Ecology 76:2135–2144.

30. Heinemeyer, K., J. Squires, M. Hebblewhite, J. J. O’Keefe, J. D. Holbrook, and J. Copeland. 2019. Wolverines in winter: indirect habitat loss and functional responses to backcountry recreation. Ecosphere 10:e02611.

31. Hilty, J., G. Worboys, A. Keeley, S. Woodley, B. Lausche, H. Locke, M. Carr, I. Pulsford, J. Pittock, and J. W. White. 2020. Guidelines for conserving connectivity through ecological networks and corridors. Gland, Switzerland: IUCN-WCPA.

32. Hooten, M. B., X. Lu, M. J. Garlick, and J. A. Powell. 2020. Animal movement models with mechanistic selection functions. Spatial Statistics 37:100406.

33. Jayadevan, A., R. Nayak, K. K. Karanth, J. Krishnaswamy, R. DeFries, K. U. Karanth, and S. Vaidyanathan. 2020. Navigating paved paradise: Evaluating landscape permeability to movement for large mammals in two conservation priority landscapes in India. Biological Conservation 247:108613.

34. Ladle, A., T. Avgar, M. Wheatley, G. B. Stenhouse, S. E. Nielsen, and M. S. Boyce. 2019. Grizzly bear response to spatio-temporal variability in human recreational activity. Journal of Applied Ecology 56:375–386.

35. Lamb, C. T., A. T. Ford, B. N. McLellan, M. F. Proctor, G. Mowat, L. Ciarniello, S. E. Nielsen, and S. Boutin. 2020. The ecology of human–carnivore coexistence. Proceedings of the National Academy of Sciences:201922097.

36. Langemann, E. G. 2011. Archaeology in the Rocky Mountain National Parks: uncovering an 11,000-year-long story. A Century of Parks Canada, 1911-2011:303-332.

37. Mahoney, P. J., G. E. Liston, S. LaPoint, E. Gurarie, B. Mangipane, A. G. Wells, T. J. Brinkman, J. U. H. Eitel, M. Hebblewhite, A. W. Nolin, N. Boelman, and L. R. Prugh. 2018. Navigating snowscapes: scale-dependent responses of mountain sheep to snowpack properties. Ecological Applications 28:1715–1729.

38. Mariela, G., C. Laura, and J. L. Belant. 2020. Planning for carnivore recolonization by mapping sex-specific landscape connectivity. Global Ecology and Conservation 21:e00869.

39. Marrotte, R. R., J. Bowman, M. G. C. Brown, C. Cordes, K. Y. Morris, M. B. Prentice, and P. J. Wilson. 2017. Multi-species genetic connectivity in a terrestrial habitat network. Movement Ecology 5:21.

40. Merkle, J. A., H. Sawyer, K. L. Monteith, S. P. H. Dwinnell, G. L. Fralick, and M. J. Kauffman. 2019. Spatial memory shapes migration and its benefits: evidence from a large herbivore. Ecology Letters 22:1797–1805.

41. Meurant, M., A. Gonzalez, A. Doxa, and C. H. Albert. 2018. Selecting surrogate species for connectivity conservation. Biological Conservation 227:326–334.

42. Michelot, T., R. Langrock, and T. A. Patterson. 2016. moveHMM: an R package for the statistical modelling of animal movement data using hidden Markov models. Methods in Ecology and Evolution 7:1308–1315.

43. Morales-González, A., H. Ruiz-Villar, A. Ordiz, and V. Penteriani. 2020. Large carnivores living alongside humans: Brown bears in human-modified landscapes. Global Ecology and Conservation 22:e00937.

44. Muff, S., J. Signer, and J. Fieberg. 2020. Accounting for individual-specific variation in habitat-selection studies: Efficient estimation of mixed-effects models using Bayesian or frequentist computation. Journal of Animal Ecology 89:80–92.

45. Naidoo, R., and A. C. Burton. 2020. Relative effects of recreational activities on a temperate terrestrial wildlife assemblage. Conservation Science and Practice 2:e271.

46. Ngoprasert, D., A. J. Lynam, and G. A. Gale. 2017. Effects of temporary closure of a national park on leopard movement and behaviour in tropical Asia. Mammalian Biology 82:65–73.

47. Nickel, B. A., J. P. Suraci, M. L. Allen, and C. C. Wilmers. 2020. Human presence and human footprint have non-equivalent effects on wildlife spatiotemporal habitat use. Biological Conservation 241:108383.

48. Nielsen, S. E., G. B. Stenhouse, and M. S. Boyce. 2006. A habitat-based framework for grizzly bear conservation in Alberta. Biological Conservation 130:217–229.

49. Palmer, S. C. F., A. Coulon, and J. M. J. Travis. 2011. Introducing a ‘stochastic movement simulator’ for estimating habitat connectivity. Methods in Ecology and Evolution 2:258–268.

50. Panzacchi, M., B. Van Moorter, O. Strand, M. Saerens, I. Kivimäki, C. C. St. Clair, I. Herfindal, and L. Boitani. 2016. Predicting the continuum between corridors and barriers to animal movements using step selection functions and randomized shortest paths. Journal of Animal Ecology 85:32–42.

51. Proctor, M. F., D. Paetkau, B. N. McLellan, G. B. Stenhouse, K. C. Kendall, R. D. Mace, W. F. Kasworm, C. Servheen, C. L. Lausen, M. L. Gibeau, W. L. Wakkinen, M. A. Haroldson, G. Mowat, C. D. Apps, L. M. Ciarniello, R. M. R. Barclay, M. S. Boyce, C. C. Schwartz, and C. Strobeck. 2012. Population fragmentation and inter-ecosystem movements of grizzly bears in western Canada and the northern United States. Wildlife Monographs 180:1–46.

52. Quaglietta, L., and M. Porto. 2019. SiMRiv: an R package for mechanistic simulation of individual, spatially-explicit multistate movements in rivers, heterogeneous and homogeneous spaces incorporating landscape bias. Movement Ecology 7:11.

53. Quaglietta, L., M. Porto, and A. T. Ford. 2019. Simulating animal movements to predict wildlife-vehicle collisions: illustrating an application of the novel R package SiMRiv. European Journal of Wildlife Research 65:100.

54. QuantumPlace Developments Ltd. 2020a. Smith Creek area structure plan, July 2020.

55. QuantumPlace Developments Ltd. 2020b. Three Sisters area structure plan, July 2020.

56. Ray, J., K. H. Redford, R. Steneck, and J. Berger. 2013. Large carnivores and the conservation of biodiversity. Island Press.

57. Ripple, W. J., J. A. Estes, R. L. Beschta, C. C. Wilmers, E. G. Ritchie, M. Hebblewhite, J. Berger, B. Elmhagen, M. Letnic, M. P. Nelson, O. J. Schmitz, D. W. Smith, A. D. Wallach, and A. J. Wirsing. 2014. Status and ecological effects of the world’s largest carnivores. Science 343.

58. Roever, C. L., H. L. Beyer, M. J. Chase, and R. J. van Aarde. 2014. The pitfalls of ignoring behaviour when quantifying habitat selection. Diversity and Distributions 20:322–333.

59. Rogala, J. T., M. Hebblewhite, J. Whittington, C. A. White, J. Coleshill, and M. Musiani. 2011. Human activity differentially redistributes large mammals in the Canadian Rockies National Parks. Ecology and Society 16:16.

60. Shepherd, B., and J. Whittington. 2006. Response of wolves in winter to wildlife corridor restoration and human use management. Ecology and Society 11:1.

61. Signer, J., J. Fieberg, and T. Avgar. 2017. Estimating utilization distributions from fitted step-selection functions. Ecosphere 8:e01771.

62. Signer, J., J. Fieberg, and T. Avgar. 2019. Animal movement tools (amt): R package for managing tracking data and conducting habitat selection analyses. Ecology and Evolution 9:880–890.

63. Suraci, J. P., L. G. Frank, A. Oriol-Cotterill, S. Ekwanga, T. M. Williams, and C. C. Wilmers. 2019. Behavior-specific habitat selection by African lions may promote their persistence in a human-dominated landscape. Ecology 100:e02644.

64. Suraci, J. P., B. A. Nickel, and C. C. Wilmers. 2020. Fine-scale movement decisions by a large carnivore inform conservation planning in human-dominated landscapes. Landscape Ecology 35:1635–1649.

65. Terborgh, J., J. A. Estes, P. Paquet, K. Ralls, D. Boyd-Herger, B. J. Miller, and R. F. Noss. 1999. The role of top carnivores in regulating terrestrial ecosystems. Pages 39-64 in M. E. Soulé and J. Terborgh, editors. Continental Conservation: Scientific Foundations of Regional Reserve Networks. The Wildlands Project. Island Pres, Washington, D.C.

66. Tucker, M. A., K. Böhning-Gaese, W. F. Fagan, J. M. Fryxell, B. Van Moorter, S. C. Alberts, A. H. Ali, A. M. Allen, N. Attias, T. Avgar, H. Bartlam-Brooks, B. Bayarbaatar, J. L. Belant, A. Bertassoni, D. Beyer, L. Bidner, F. M. van Beest, S. Blake, N. Blaum, C. Bracis, D. Brown, P. J. N. de Bruyn, F. Cagnacci, J. M. Calabrese, C. Camilo-Alves, S. Chamaillé-Jammes, A. Chiaradia, S. C. Davidson, T. Dennis, S. DeStefano, D. Diefenbach, I. Douglas-Hamilton, J. Fennessy, C. Fichtel, W. Fiedler, C. Fischer, I. Fischhoff, C. H. Fleming, A. T. Ford, S. A. Fritz, B. Gehr, J. R. Goheen, E. Gurarie, M. Hebblewhite, M. Heurich, A. J. M. Hewison, C. Hof, E. Hurme, L. A. Isbell, R. Janssen, F. Jeltsch, P. Kaczensky, A. Kane, P. M. Kappeler, M. Kauffman, R. Kays, D. Kimuyu, F. Koch, B. Kranstauber, S. LaPoint, P. Leimgruber, J. D. C. Linnell, P. López-López, A. C. Markham, J. Mattisson, E. P. Medici, U. Mellone, E. Merrill, G. de Miranda Mourão, R. G. Morato, N. Morellet, T. A. Morrison, S. L. Díaz-Muñoz, A. Mysterud, D. Nandintsetseg, R. Nathan, A. Niamir, J. Odden, R. B. O’Hara, L. G. R. Oliveira-Santos, K. A. Olson, B. D. Patterson, R. Cunha de Paula, L. Pedrotti, B. Reineking, M. Rimmler, T. L. Rogers, C. M. Rolandsen, C. S. Rosenberry, D. I. Rubenstein, K. Safi, S. Saïd, N. Sapir, H. Sawyer, N. M. Schmidt, N. Selva, A. Sergiel, E. Shiilegdamba, J. P. Silva, N. Singh, E. J. Solberg, O. Spiegel, O. Strand, S. Sundaresan, W. Ullmann, U. Voigt, J. Wall, D. Wattles, M. Wikelski, C. C. Wilmers, J. W. Wilson, G. Wittemyer, F. Zięba, T. Zwijacz-Kozica, and T. Mueller. 2018. Moving in the Anthropocene: Global reductions in terrestrial mammalian movements. Science 359:466–469.

67. Wang, F., W. J. McShea, D. Wang, S. Li, Q. Zhao, H. Wang, and Z. Lu. 2014. Evaluating landscape options for corridor restoration between giant panda reserves. PLoS ONE 9:e105086.

68. Whittington, J., P. Low, and B. Hunt. 2019. Temporal road closures improve habitat quality for wildlife. Scientific reports 9:3772.

69. Whittington, J., C. C. St. Clair, and G. Mercer. 2005. Spatial responses of wolves to roads and trails in mountain valleys. Ecological Applications 15:543–553.

70. Wittemyer, G., P. Elsen, W. T. Bean, A. C. O. Burton, and J. S. Brashares. 2008. Accelerated human population growth at protected area edges. Science 321:123–126.

71. Zeller, K., K. McGarigal, S. Cushman, P. Beier, T. W. Vickers, and W. Boyce. 2015. Using step and path selection functions for estimating resistance to movement: pumas as a case study. Landscape Ecology:1–17.

72. Zeller, K. A., M. K. Jennings, T. W. Vickers, H. B. Ernest, S. A. Cushman, and W. M. Boyce. 2018. Are all data types and connectivity models created equal? Validating common connectivity approaches with dispersal data. Diversity and Distributions 24:868–879.

73. Zeller, K. A., D. W. Wattles, J. M. Bauder, and S. DeStefano. 2020. Forecasting seasonal habitat connectivity in a developing landscape. Land 9:233.

74. Zeller, K. A., D. W. Wattles, L. Conlee, and S. DeStefano. 2019. Black bears alter movements in response to anthropogenic features with time of day and season. Movement Ecology 7:19.

75. Zhai, Y., P. Korça Baran, and C. Wu. 2018. Can trail spatial attributes predict trail use level in urban forest park? An examination integrating GPS data and space syntax theory. Urban Forestry & Urban Greening 29:171–182.

