## Appendix S1 for "Towns and Trails Drive Carnivore Connectivity using a Step Selection Approach"

### Appendix S1. Maps of study area, observed GPS locations, and predicted utilization distributions from step selection function models.

#### *Section S1.1 Study area and GPS locations*

Research Permits: Researchers fit wolves and grizzly bears with GPS collars for several research projects with the following permits: University of Alberta Animal care protocol ID# 353112, University of Montana Institutional Animal Care Protocol 059-08MHECS-120908, 004-16MHECS-020916, 066-18MHWB-123118, Parks Canada Research and Collection Permits LL-2010-4392, BAN-2015-18276, BAN-2018-30898, LL- 2012-10975.

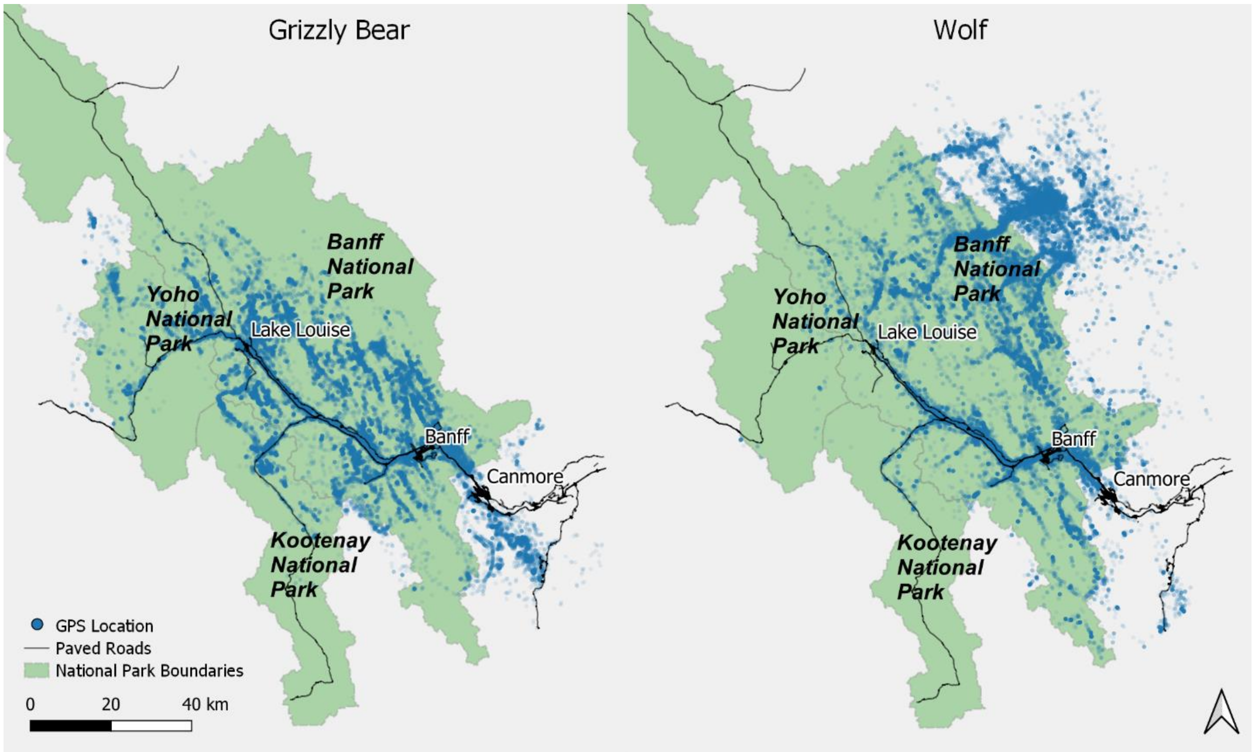


Figure S1. Grizzly bear and wolf GPS locations within and adjacent to Banff, Kootenay, and Yoho National Parks of Canada, 2000 to 2020. We used GPS locations from this study area to develop step selection and resource selection function models.

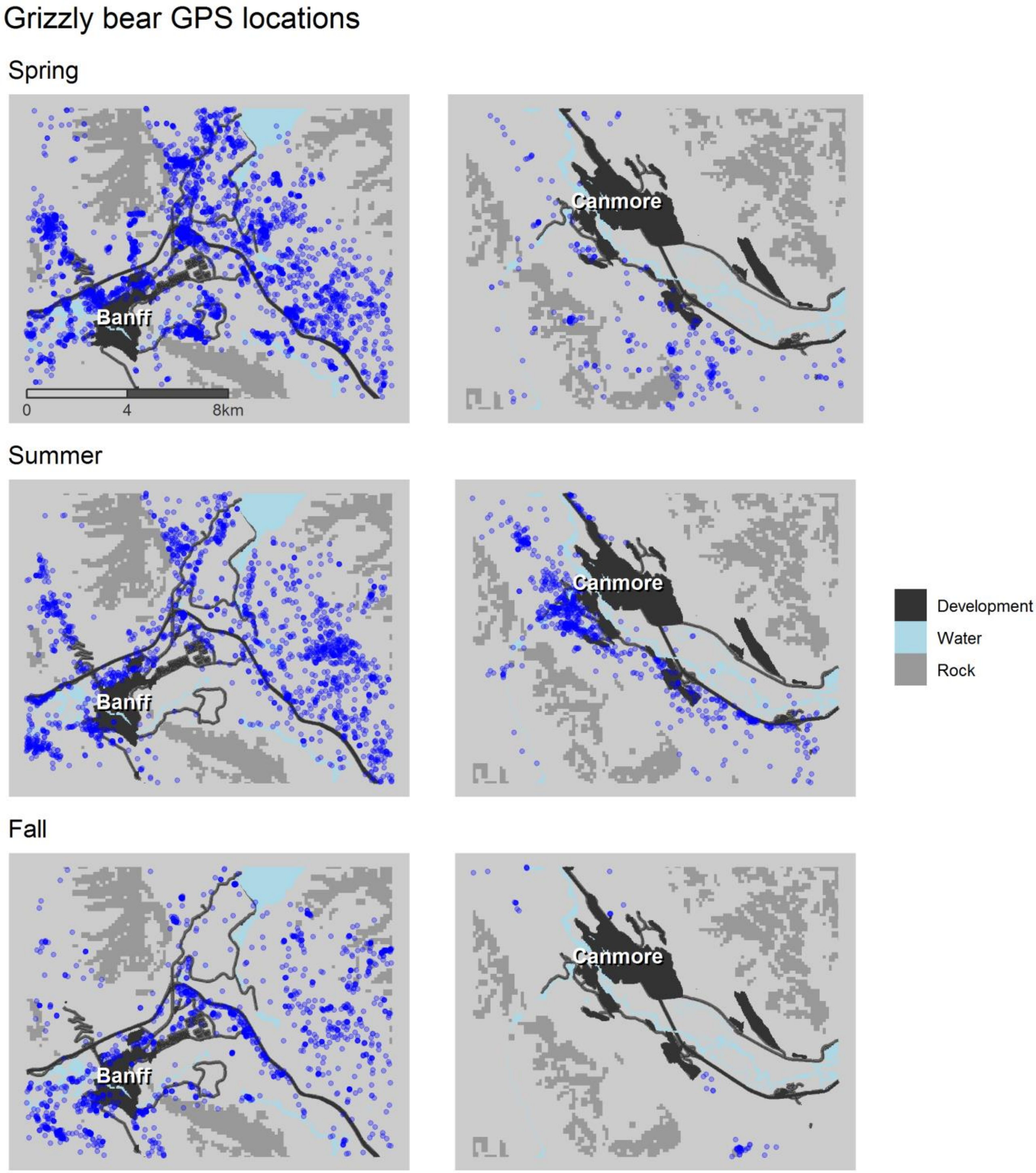


Figure S2. Distribution of grizzly bear locations around the towns of Banff and Canmore by season.

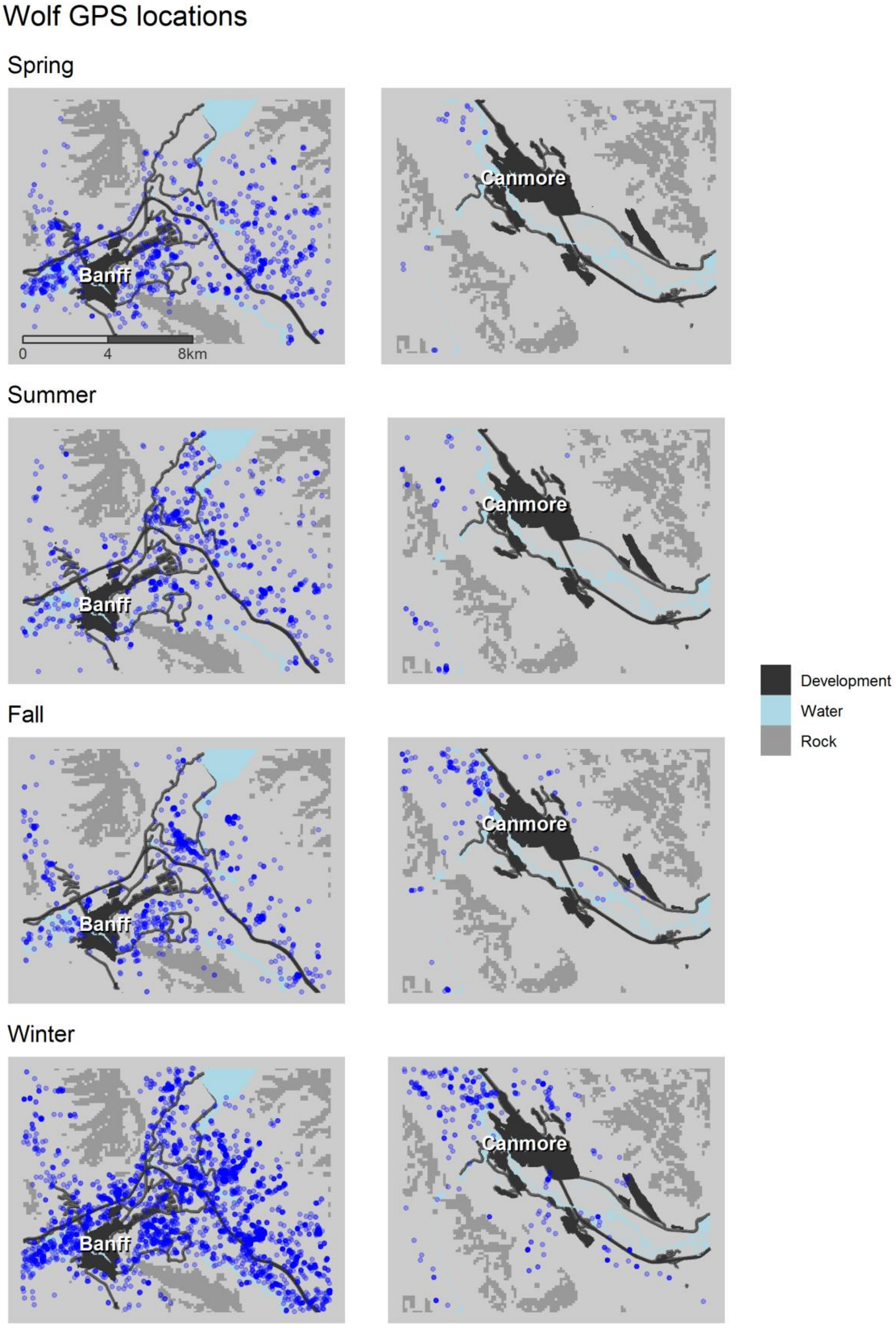


Figure S3. Distribution of wolf locations around the towns of Banff and Canmore by season.

*Section S1.2 Grizzly bear utilization distributions and change in utilization distribution by season in the Bow Valley*

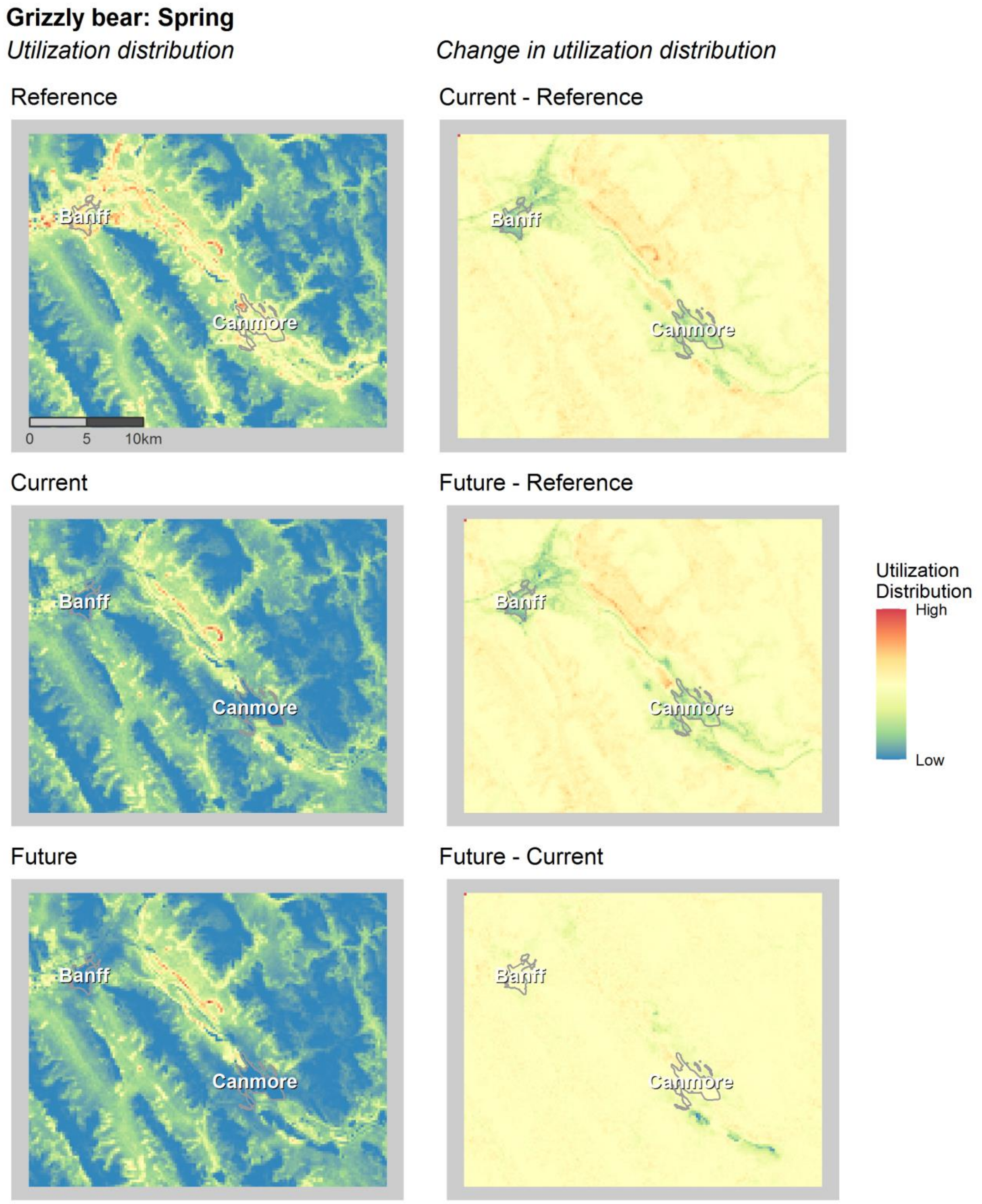

**Grizzly bear: Summer**

*Utilization distribution*

*Change in utilization distribution*

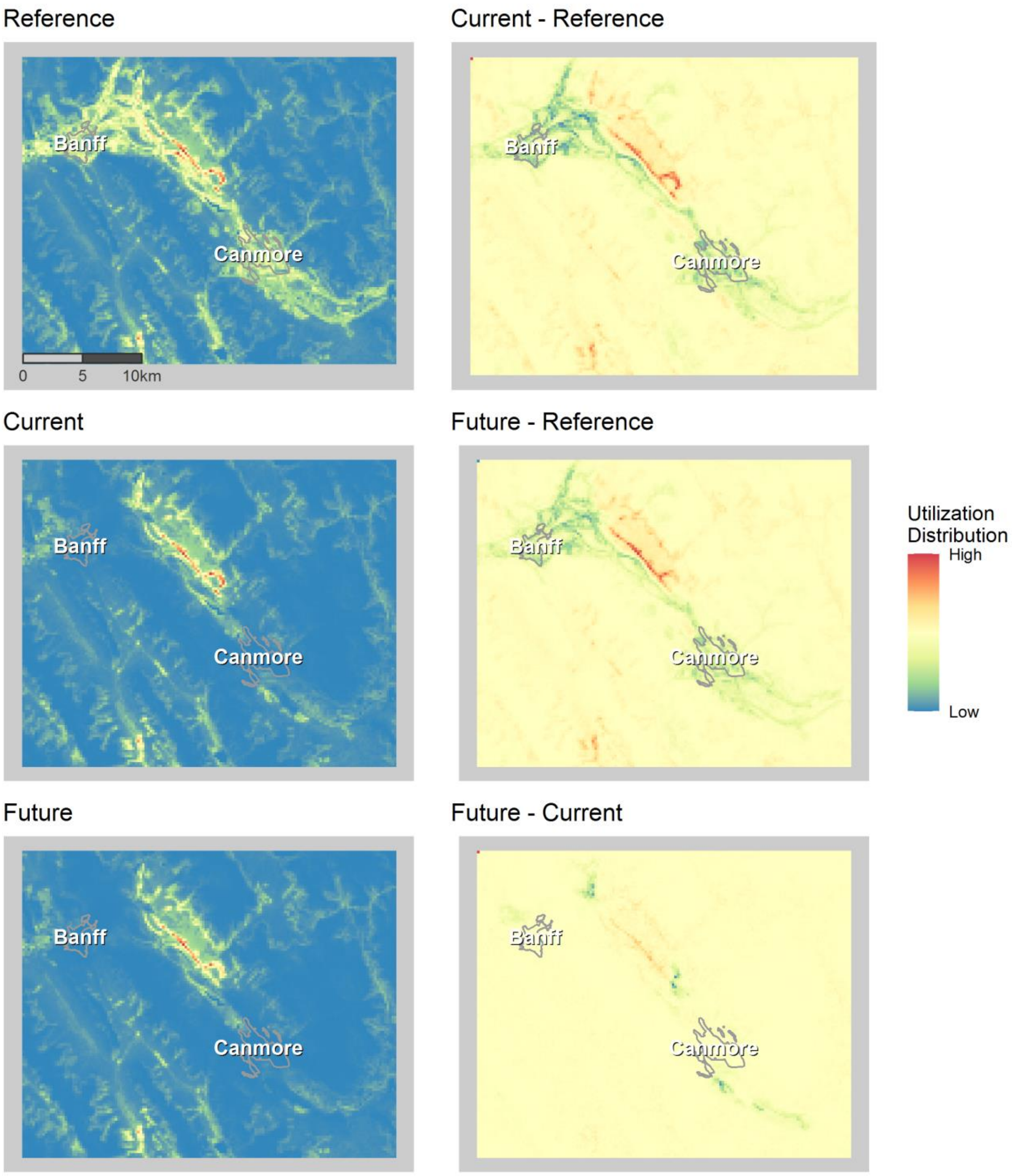

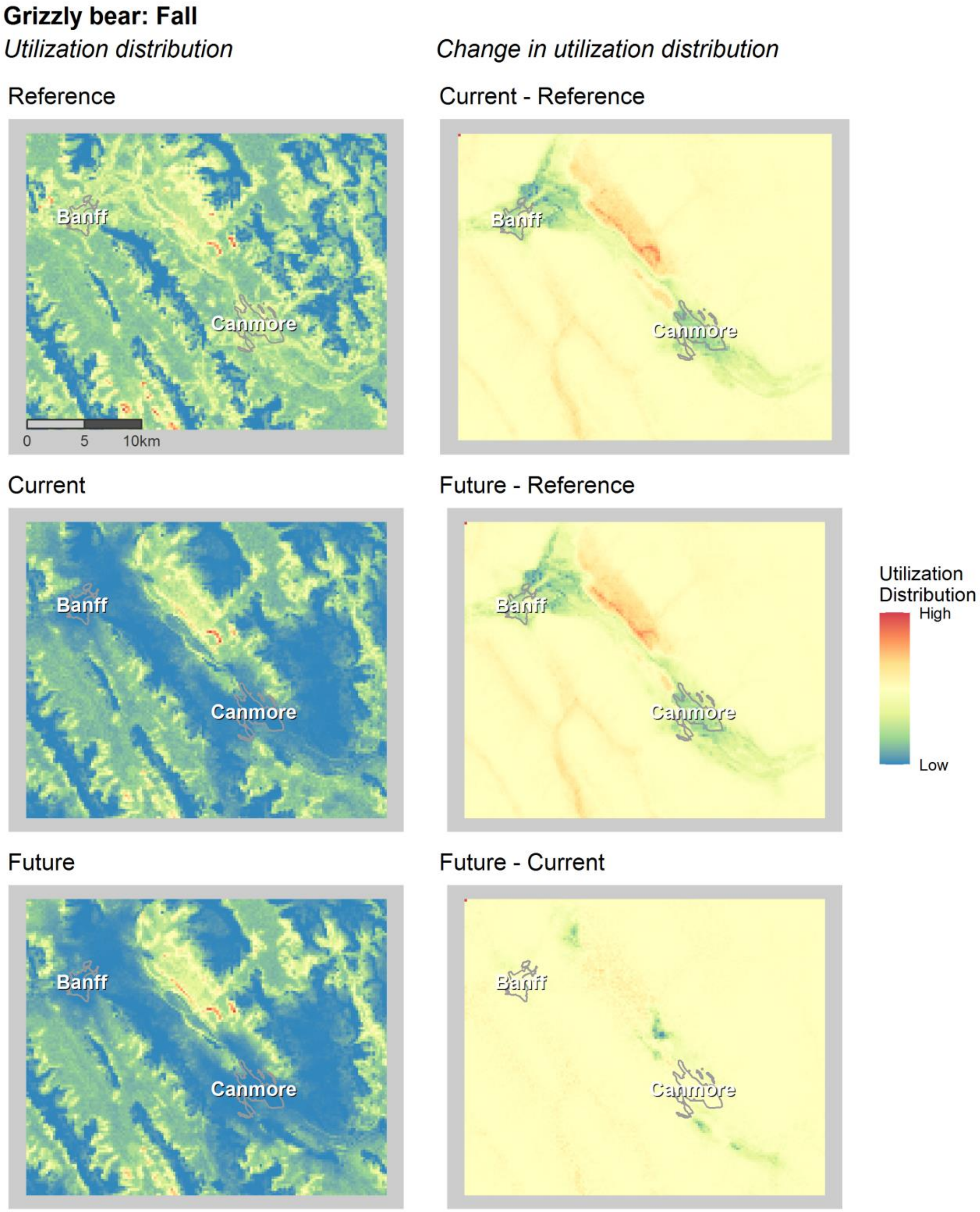


Figure S4. Grizzly bear utilization distributions (left) and changes in utilization distributions between time periods (right) near the towns of Banff and Canmore.

*Section S1.3 Wolf utilization distributions and change in utilization distribution by season in the Bow Valley*

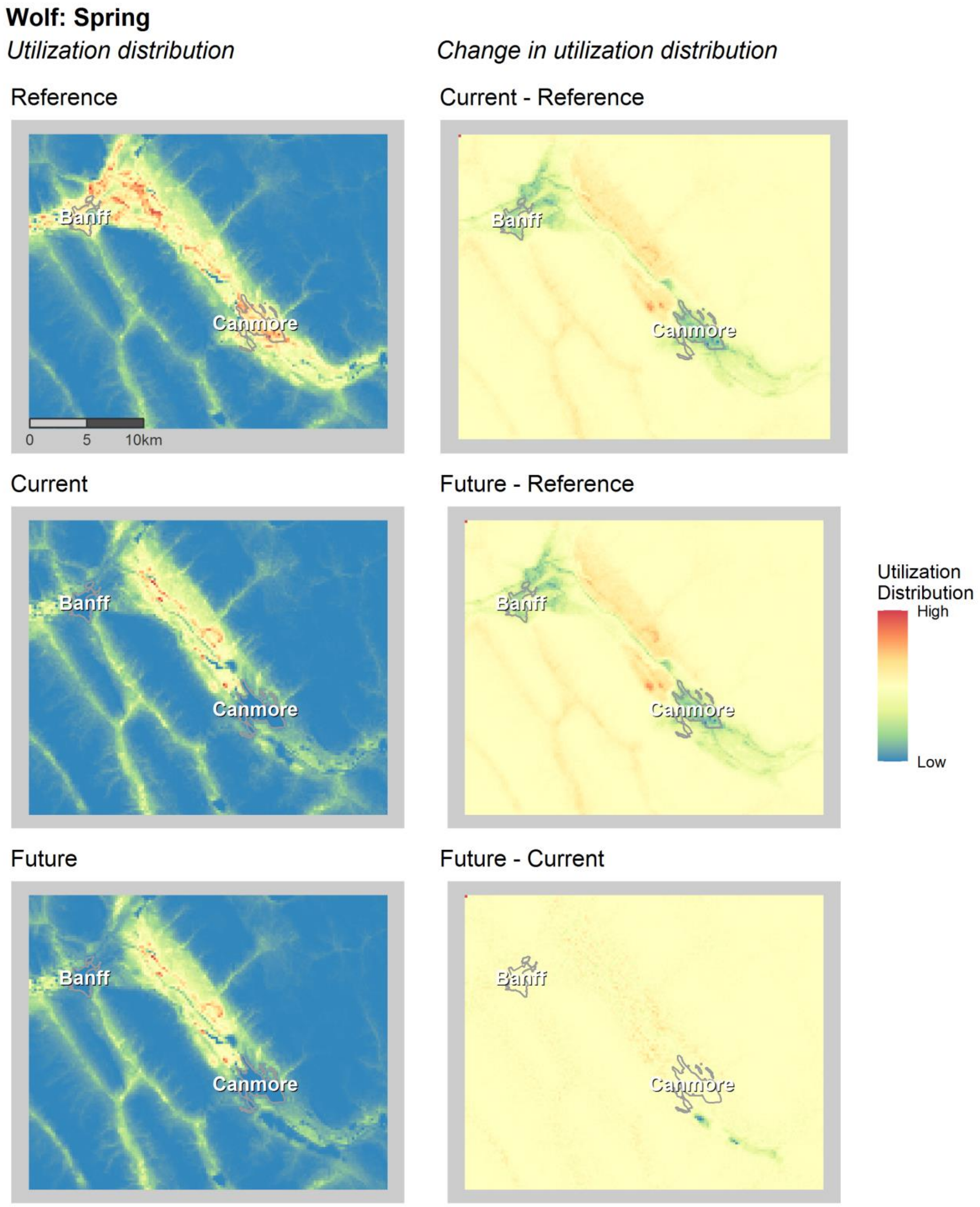

### Wolf: Summer

*Utilization distribution*

*Change in utilization distribution*

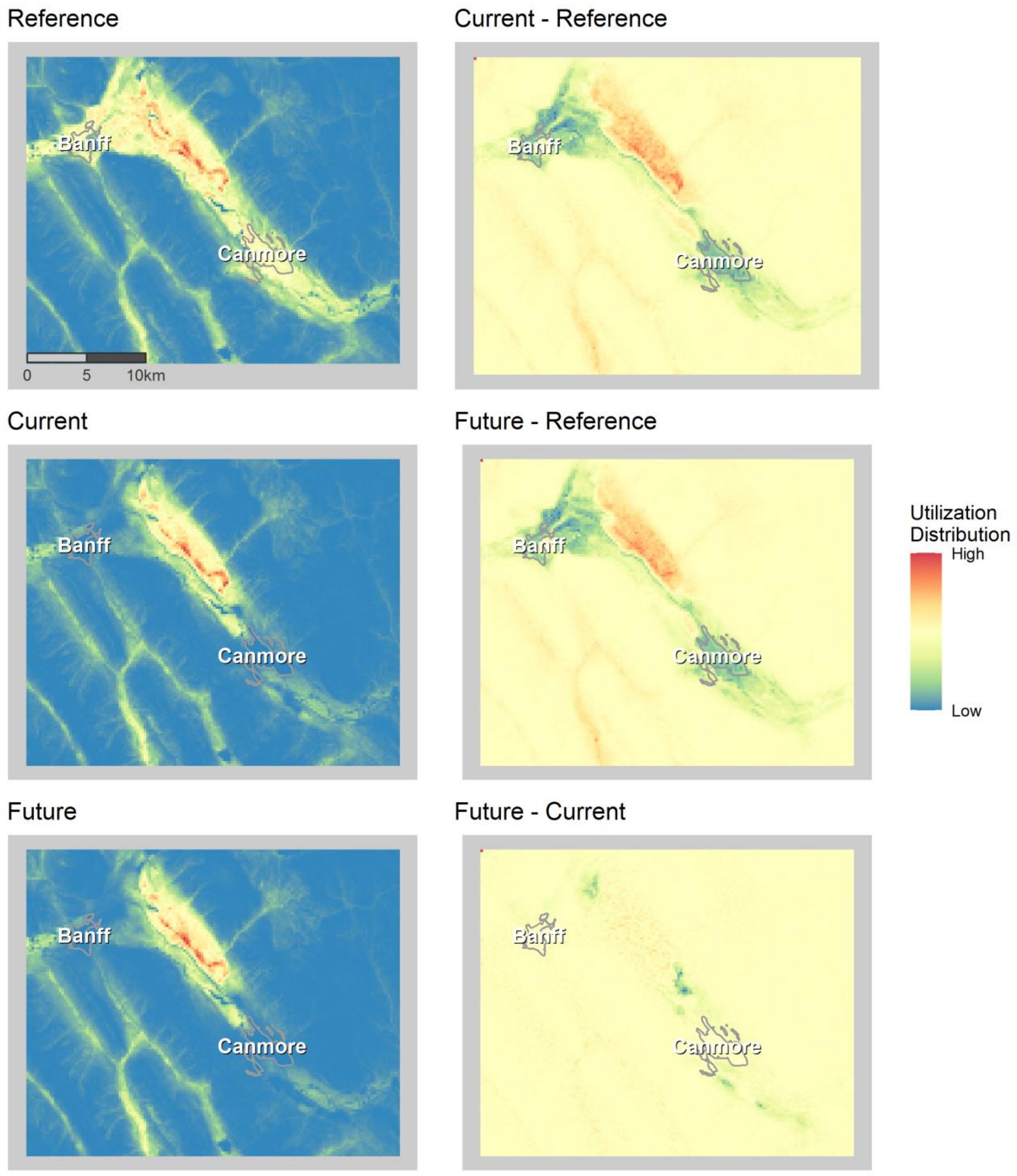

**Wolf: Fall**

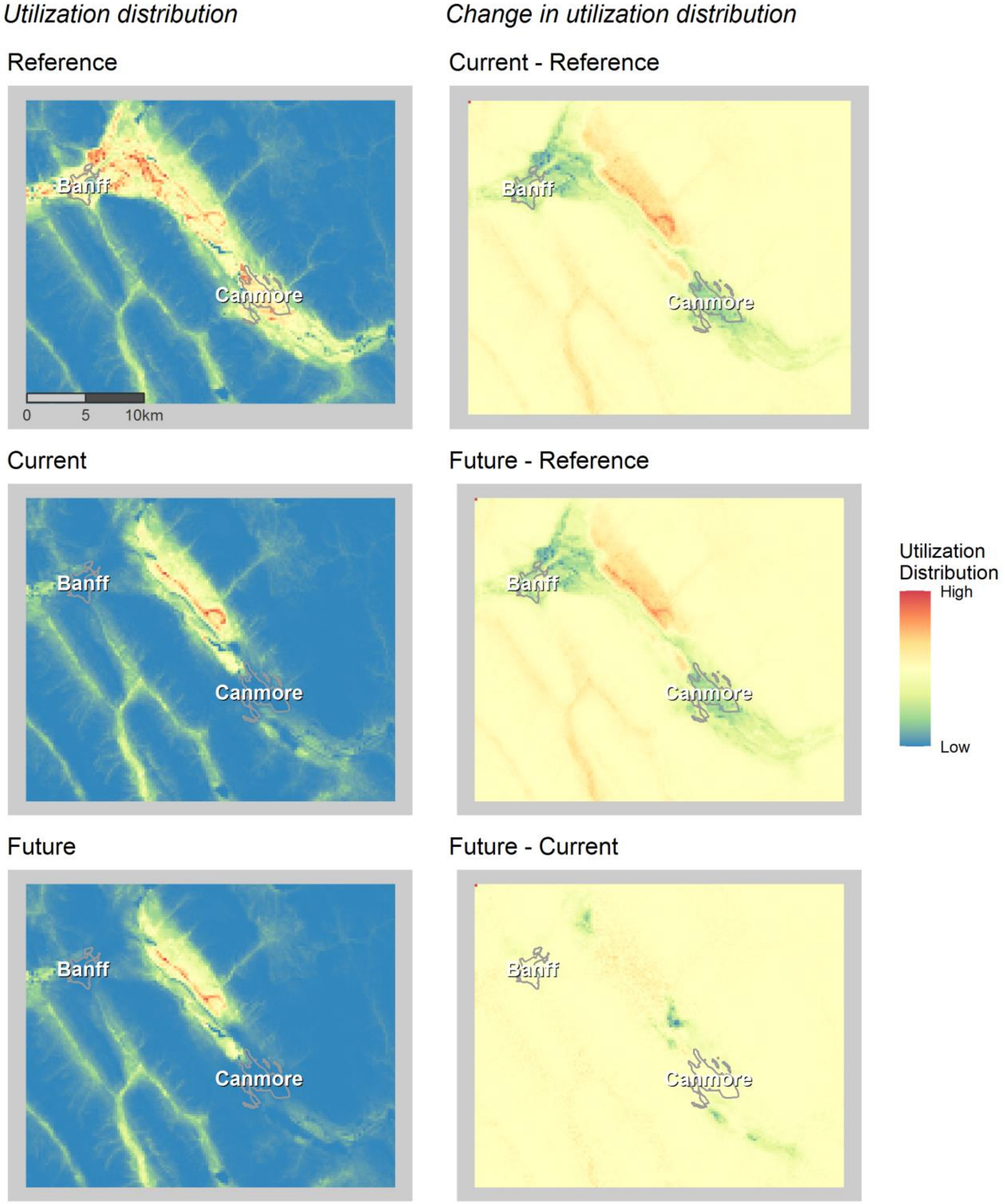

Utilization distribution
Change in utilization distribution
Reference
Banff
Canmore
0
5
10km
Current - Reference
Banff
Canmore
Current
Banff
Canmore
Future - Reference
Banff
Canmore
Utilization Distribution
High
Low
Future
Banff
Canmore
Future - Current
Banff
Canmore

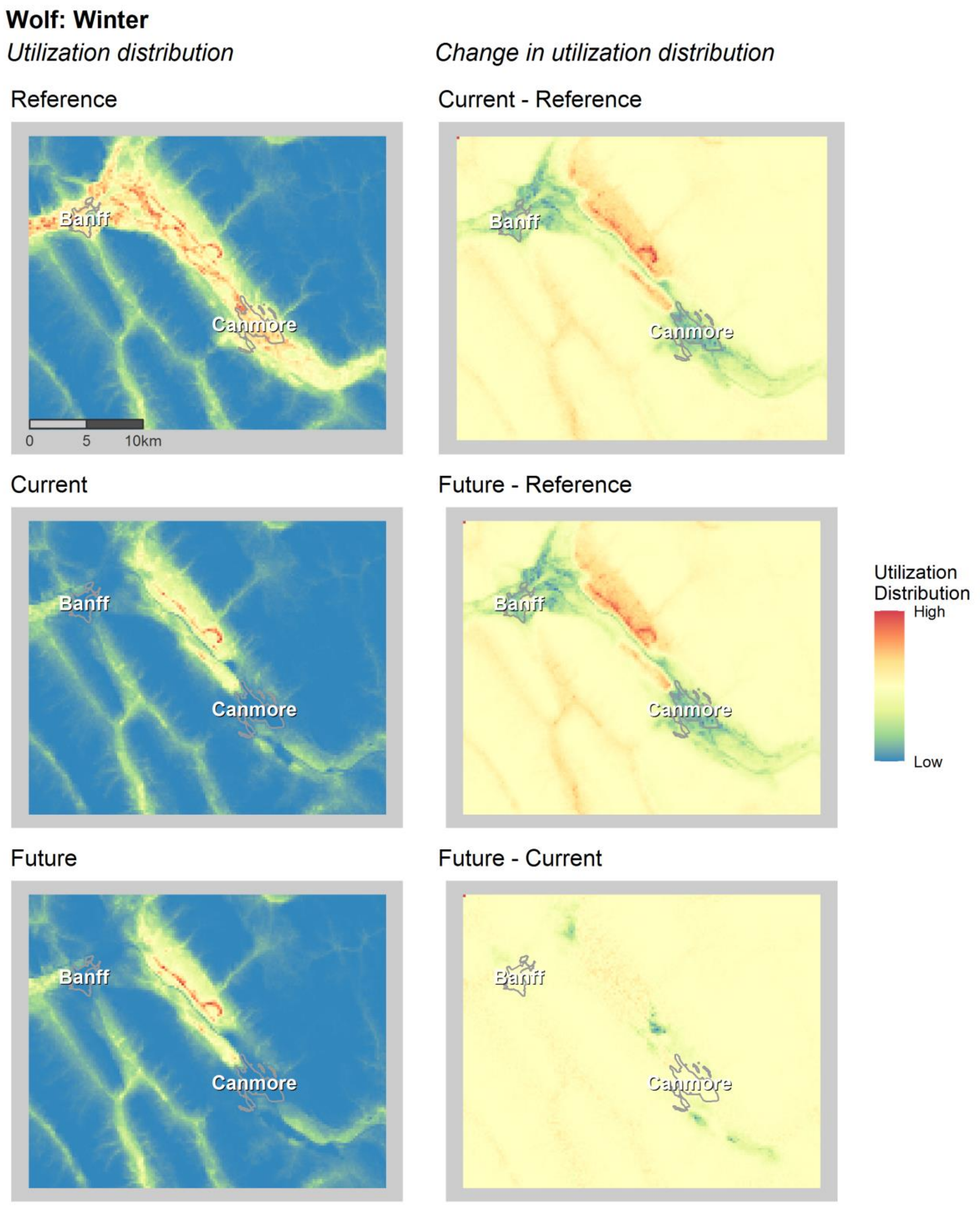


Figure S5. Wolf utilization distributions (left) and changes in utilization distribution (right) near the towns of Banff and Canmore.
