## Appendix S2 for "Towns and Trails Drive Carnivore Connectivity using a Step Selection Approach"

### Appendix S2: Tables of parameter estimates from movement models, step selection functions, and resource selection functions.

#### *Section 2.1 Explanatory variables*

Table S1. Explanatory variables used and considered in wolf and grizzly bear step selection and resource selection analyses. We standardized most continuous variables by their mean and standard deviation to improve convergence and interpretability. Otherwise, we applied a decay function to most distance covariates so that the effect of the covariate declined with distance with an asymptote occurring around 500 m.

| **Explanatory Variable** | **Transformation** |
|---|---|
| *INCLUDED IN FINAL MODELS* | |
| Distance to campground (m) | 1 – exp(-10 * x * 0.001) |
| Distance to town (m) | 1 – exp(-10 * x * 0.001) |
| Trail density (km $km^{-2}$) formal trails, 500 m radius | (x – mean(x)) / 10*sd(x) |
| Trail density (km $km^{-2}$) x ln(Distance to paved road (km)) | |
| Off/on railway (0 = off, 1 = on) | |
| Off/on trail (0 = off, 1 = on) | |
| Elevation (m) | (x – mean(x)) / sd(x) |
| Slope (degrees) | (x – mean(x)) / sd(x) |
| Aspect South [-1 * cosine(aspect)]; (1 = south, -1 = north, 0 = east or west) | (x – mean(x)) / sd(x) |
| Land Cover | |
| Forest – closed coniferous (reference category*)* | |
| Forest – deciduous and coniferous | |
| Herbaceous | |
| Shrub | |
| Barren and rock | |
| Distance to edge of forest (m) | 1 – exp(-10 * x * 0.001) |
| Distance (m) to large vegetated patch $\geq$ 9 $km^2$ | (x – mean(x)) / sd(x) |
| Distance to stream (m) | 1 – exp(-10 * x * 0.001) |
| Burned (0 = no, 1 = forest burned since 1960) | |
| *EXCLUDED FROM FINAL MODELS* | |
| Trail density (km $km^{-2}$) for all formal and informal trails, 500 m radius | (x – mean(x)) / sd(x) |
| Normalized difference vegetation index (NDVI) (July and August), mean and standard deviation | (x – mean(x)) / sd(x) |
| Dynamic habitat index (DHI) – cumulative NDVI | (x – mean(x)) / sd(x) |
| Percent crown cover | (x – mean(x)) / sd(x) |
| Seasonal mean decadal temperature (degrees C) from ClimateNA | (x – mean(x)) / sd(x) |
| Seasonal mean decadal precipitation (mm) from ClimateNA | (x – mean(x)) / sd(x) |

### *Section 2.2 Hidden Markov movement model results*

Table S2. Parameter estimates from hidden Markov movement models. The movement models contained two movement states (slow and fast) and state specific parameters for step length (gamma distribution) and turn angle (von Mises distribution). The gamma distribution contained the parameters mean and standard deviation (SD). The link to from mean and SD to commonly used shape and rate is given by: shape = $mean^2/SD^2$ and rate = $mean/SD^2$. The von Mises distribution for turn angles included mean and concentration. Movement state transitions depended on the time of day such that Cosine Hour = cosine(Hour * π/12).

| ***Species*** | ***Season*** | ***Type*** | ***Parameter*** | ***State*** | ***Estimate*** | ***LCL*** | ***UCL*** |
|---|---|---|---|---|---|---|---|
| Grizzly Bear | Fall | Movement Parameter | Step Length Mean | Slow | 0.012 | 0.011 | 0.011 |
| | | Movement Parameter | Step Length Mean | Fast | 0.643 | 0.629 | 0.743 |
| | | Movement Parameter | Step Length SD | Slow | 0.010 | 0.010 | 0.011 |
| | | Movement Parameter | Step Length SD | Fast | 0.726 | 0.709 | 0.743 |
| | | Movement Parameter | Turn Angle Mean | Slow | 3.124 | 3.062 | 0.717 |
| | | Movement Parameter | Turn Angle Mean | Fast | -0.017 | -0.068 | 0.543 |
| | | Movement Parameter | Turn Angle Concentration | Slow | 0.672 | 0.628 | 0.717 |
| | | Movement Parameter | Turn Angle Concentration | Fast | 0.516 | 0.488 | 0.543 |
| | | Transition Coefficient | Intercept | Slow to Fast | -0.572 | -0.650 | -0.639 |
| | | Transition Coefficient | Intercept | Fast to Slow | -2.689 | -2.835 | 4.088 |
| | | Transition Coefficient | Cosine Hour | Slow to Fast | -0.748 | -0.858 | -0.639 |
| | | Transition Coefficient | Cosine Hour | Fast to Slow | 3.851 | 3.615 | 4.088 |

| ***Species*** | ***Season*** | ***Type*** | ***Parameter*** | ***State*** | ***Estimate*** | ***LCL*** | ***UCL*** |
|---|---|---|---|---|---|---|---|
| Grizzly Bear | Spring | Movement Parameter | Step Length Mean | Slow | 0.121 | 0.115 | 0.161 |
| | | Movement Parameter | Step Length Mean | Fast | 1.425 | 1.374 | 1.493 |
| | | Movement Parameter | Step Length SD | Slow | 0.152 | 0.144 | 0.161 |
| | | Movement Parameter | Step Length SD | Fast | 1.449 | 1.407 | 1.493 |
| | | Movement Parameter | Turn Angle Mean | Slow | 3.106 | 3.024 | 0.368 |
| | | Movement Parameter | Turn Angle Mean | Fast | -0.003 | -0.044 | 0.953 |
| | | Movement Parameter | Turn Angle Concentration | Slow | 0.336 | 0.305 | 0.368 |
| | | Movement Parameter | Turn Angle Concentration | Fast | 0.902 | 0.851 | 0.953 |
| | | Transition Coefficient | Intercept | Slow to Fast | -1.785 | -1.857 | -0.034 |
| | | Transition Coefficient | Intercept | Fast to Slow | -1.248 | -1.339 | 1.140 |
| | | Transition Coefficient | Cosine Hour | Slow to Fast | -0.123 | -0.211 | -0.034 |
| | | Transition Coefficient | Cosine Hour | Fast to Slow | 1.011 | 0.882 | 1.140 |
| Grizzly Bear | Summer | Movement Parameter | Step Length Mean | Slow | 0.013 | 0.013 | 0.012 |
| | | Movement Parameter | Step Length Mean | Fast | 0.801 | 0.789 | 0.919 |
| | | Movement Parameter | Step Length SD | Slow | 0.011 | 0.011 | 0.012 |

| ***Species*** | ***Season*** | ***Type*** | ***Parameter*** | ***State*** | ***Estimate*** | ***LCL*** | ***UCL*** |
|---|---|---|---|---|---|---|---|
| | | Movement Parameter | Step Length SD | Fast | 0.903 | 0.887 | 0.919 |
| | | Movement Parameter | Turn Angle Mean | Slow | 3.140 | 3.057 | 0.582 |
| | | Movement Parameter | Turn Angle Mean | Fast | -0.012 | -0.053 | 0.495 |
| | | Movement Parameter | Turn Angle Concentration | Slow | 0.535 | 0.489 | 0.582 |
| | | Movement Parameter | Turn Angle Concentration | Fast | 0.474 | 0.454 | 0.495 |
| | | Transition Coefficient | Intercept | Slow to Fast | 11.739 | 10.468 | - 13.547 |
| | | Transition Coefficient | Intercept | Fast to Slow | -7.963 | -8.503 | 9.990 |
| | | Transition Coefficient | Cosine Hour | Slow to Fast | -15.046 | - 16.544 | - 13.547 |
| | | Transition Coefficient | Cosine Hour | Fast to Slow | 9.352 | 8.714 | 9.990 |
| Wolf | Fall | Movement Parameter | Step Length Mean | Slow | 0.025 | 0.024 | 0.028 |
| | | Movement Parameter | Step Length Mean | Fast | 2.185 | 2.109 | 2.743 |
| | | Movement Parameter | Step Length SD | Slow | 0.026 | 0.024 | 0.028 |
| | | Movement Parameter | Step Length SD | Fast | 2.650 | 2.560 | 2.743 |
| | | Movement Parameter | Turn Angle Mean | Slow | -3.111 | -3.223 | 0.478 |
| | | Movement Parameter | Turn Angle Mean | Fast | 0.026 | -0.100 | 0.337 |

| ***Species*** | ***Season*** | ***Type*** | ***Parameter*** | ***State*** | ***Estimate*** | ***LCL*** | ***UCL*** |
|---|---|---|---|---|---|---|---|
| | | Movement Parameter | Turn Angle Concentration | Slow | 0.427 | 0.376 | 0.478 |
| | | Movement Parameter | Turn Angle Concentration | Fast | 0.297 | 0.259 | 0.337 |
| | | Transition Coefficient | Intercept | Slow to Fast | -0.652 | -0.730 | -0.196 |
| | | Transition Coefficient | Intercept | Fast to Slow | -1.275 | -1.358 | 0.005 |
| | | Transition Coefficient | Cosine Hour | Slow to Fast | -0.306 | -0.417 | -0.196 |
| | | Transition Coefficient | Cosine Hour | Fast to Slow | -0.100 | -0.205 | 0.005 |
| Wolf | Spring | Movement Parameter | Step Length Mean | Slow | 0.034 | 0.033 | 0.035 |
| | | Movement Parameter | Step Length Mean | Fast | 2.456 | 2.384 | 3.129 |
| | | Movement Parameter | Step Length SD | Slow | 0.033 | 0.031 | 0.035 |
| | | Movement Parameter | Step Length SD | Fast | 3.038 | 2.950 | 3.129 |
| | | Movement Parameter | Turn Angle Mean | Slow | 3.065 | 2.996 | 0.588 |
| | | Movement Parameter | Turn Angle Mean | Fast | -0.028 | -0.110 | 0.417 |
| | | Movement Parameter | Turn Angle Concentration | Slow | 0.546 | 0.504 | 0.588 |
| | | Movement Parameter | Turn Angle Concentration | Fast | 0.384 | 0.351 | 0.417 |
| | | Transition Coefficient | Intercept | Slow to Fast | -0.867 | -0.930 | 0.264 |

| *Species* | *Season* | *Type* | *Parameter* | *State* | *Estimate* | *LCL* | *UCL* |
|---|---|---|---|---|---|---|---|
| | | Transition Coefficient | Intercept | Fast to Slow | -1.312 | -1.384 | -0.214 |
| | | Transition Coefficient | Cosine Hour | Slow to Fast | 0.176 | 0.087 | 0.264 |
| | | Transition Coefficient | Cosine Hour | Fast to Slow | -0.305 | -0.395 | -0.214 |
| Wolf | Summer | Movement Parameter | Step Length Mean | Slow | 0.036 | 0.033 | 0.041 |
| | | Movement Parameter | Step Length Mean | Fast | 2.242 | 2.176 | 2.807 |
| | | Movement Parameter | Step Length SD | Slow | 0.038 | 0.035 | 0.041 |
| | | Movement Parameter | Step Length SD | Fast | 2.730 | 2.655 | 2.807 |
| | | Movement Parameter | Turn Angle Mean | Slow | 3.107 | 3.016 | 0.516 |
| | | Movement Parameter | Turn Angle Mean | Fast | 0.055 | -0.039 | 0.356 |
| | | Movement Parameter | Turn Angle Concentration | Slow | 0.468 | 0.421 | 0.516 |
| | | Movement Parameter | Turn Angle Concentration | Fast | 0.324 | 0.292 | 0.356 |
| | | Transition Coefficient | Intercept | Slow to Fast | -0.501 | -0.569 | 0.047 |
| | | Transition Coefficient | Intercept | Fast to Slow | -1.353 | -1.428 | -0.097 |
| | | Transition Coefficient | Cosine Hour | Slow to Fast | -0.049 | -0.145 | 0.047 |
| | | Transition Coefficient | Cosine Hour | Fast to Slow | -0.186 | -0.276 | -0.097 |

| ***Species*** | ***Season*** | ***Type*** | ***Parameter*** | ***State*** | ***Estimate*** | ***LCL*** | ***UCL*** |
|---|---|---|---|---|---|---|---|
| Wolf | Winter | Movement Parameter | Step Length Mean | Slow | 0.023 | 0.022 | 0.023 |
| | | Movement Parameter | Step Length Mean | Fast | 1.824 | 1.791 | 2.253 |
| | | Movement Parameter | Step Length SD | Slow | 0.022 | 0.021 | 0.023 |
| | | Movement Parameter | Step Length SD | Fast | 2.213 | 2.173 | 2.253 |
| | | Movement Parameter | Turn Angle Mean | Slow | 3.081 | 3.035 | 0.558 |
| | | Movement Parameter | Turn Angle Mean | Fast | -0.008 | -0.054 | 0.449 |
| | | Movement Parameter | Turn Angle Concentration | Slow | 0.531 | 0.504 | 0.558 |
| | | Movement Parameter | Turn Angle Concentration | Fast | 0.428 | 0.407 | 0.449 |
| | | Transition Coefficient | Intercept | Slow to Fast | -0.695 | -0.734 | -0.246 |
| | | Transition Coefficient | Intercept | Fast to Slow | -1.212 | -1.254 | -0.138 |
| | | Transition Coefficient | Cosine Hour | Slow to Fast | -0.302 | -0.358 | -0.246 |
| | | Transition Coefficient | Cosine Hour | Fast to Slow | -0.192 | -0.245 | -0.138 |

### *Section 2.3 Step selection function results*

Table S3. SSF parameter estimates and 95% confidence intervals for grizzly bears and wolves by season in Banff National Park and surrounding areas, 2002 - 2020.

| *Species* | *Season* | *Parameter* | *Estimate* | *SE* | *Wald* | *p-value* | *LCL* | *UCL* |
|---|---|---|---|---|---|---|---|---|
| Grizzly Bear | Fall | Burned since 1960 | 0.44 | 0.07 | 6.7 | <0.001 | 0.31 | 0.57 |
| | | Cosine Turn Angle | -2.49 | 0.05 | -54.2 | <0.001 | -2.58 | -2.40 |
| | | Cosine Turn Angle:FastState | 3.37 | 0.06 | 55.9 | <0.001 | 3.25 | 3.49 |
| | | Distance Campground | 1.08 | 0.43 | 2.5 | 0.012 | 0.23 | 1.92 |
| | | Distance to Forest Edge | -0.86 | 0.20 | -4.2 | <0.001 | -1.26 | -0.46 |
| | | Distance to Patch > 9 km2 | -1.20 | 0.05 | -22.3 | <0.001 | -1.31 | -1.10 |
| | | Distance Town | 0.44 | 0.26 | 1.7 | 0.089 | -0.07 | 0.94 |
| | | Distance Town:Night | -0.07 | 0.01 | -5.0 | <0.001 | -0.09 | -0.04 |
| | | Elevation | -0.05 | 0.17 | -0.3 | 0.779 | -0.39 | 0.29 |
| | | Landcover: Barren | 0.28 | 0.14 | 2.0 | 0.049 | 0.00 | 0.55 |
| | | Landcover: Herbaceous | 0.96 | 0.16 | 6.2 | <0.001 | 0.65 | 1.26 |
| | | Landcover: Open Conifer or Deciduous | 0.30 | 0.08 | 3.6 | <0.001 | 0.14 | 0.47 |
| | | Landcover: Shrub | 0.59 | 0.12 | 4.9 | <0.001 | 0.35 | 0.82 |
| | | logStepLength | -0.90 | 0.01 | -87.7 | <0.001 | -0.92 | -0.88 |
| | | logStepLength:FastState | 1.17 | 0.01 | 102.7 | <0.001 | 1.15 | 1.19 |
| | | Railway | 1.62 | 0.12 | 13.8 | <0.001 | 1.39 | 1.85 |
| | | Slope | 0.08 | 0.08 | 1.1 | 0.285 | -0.07 | 0.23 |
| | | Southerly Aspect | 0.04 | 0.05 | 0.9 | 0.387 | -0.05 | 0.13 |

| *Species* | *Season* | *Parameter* | *Estimate* | *SE* | *Wald* | *p-value* | *LCL* | *UCL* |
|---|---|---|---|---|---|---|---|---|
| | | Trail | 0.05 | 0.19 | 0.3 | 0.799 | -0.32 | 0.42 |
| | | TrailDensity | -1.32 | 0.51 | -2.6 | 0.009 | -2.32 | -0.32 |
| | | TrailDensity:logDistPavedRoad | 0.02 | 0.01 | 2.3 | 0.023 | 0.00 | 0.05 |
| Grizzly Bear | Spring | Burned since 1960 | 0.07 | 0.05 | 1.3 | 0.208 | -0.04 | 0.17 |
| | | Cosine Turn Angle | -1.02 | 0.02 | -45.6 | <0.001 | -1.06 | -0.98 |
| | | Cosine Turn Angle:FastState | 2.01 | 0.04 | 49.8 | <0.001 | 1.93 | 2.09 |
| | | Distance Campground | 0.21 | 0.15 | 1.4 | 0.167 | -0.09 | 0.51 |
| | | Distance to Forest Edge | -0.99 | 0.11 | -9.3 | <0.001 | -1.20 | -0.78 |
| | | Distance to Patch > 9 km2 | -0.55 | 0.06 | -9.4 | <0.001 | -0.66 | -0.43 |
| | | Distance Town | 0.61 | 0.14 | 4.2 | <0.001 | 0.32 | 0.89 |
| | | Distance Town:Night | -0.03 | 0.01 | -2.8 | 0.006 | -0.05 | -0.01 |
| | | Elevation | -0.78 | 0.15 | -5.1 | <0.001 | -1.07 | -0.48 |
| | | Landcover: Barren | 0.17 | 0.09 | 1.9 | 0.053 | -0.00 | 0.35 |
| | | Landcover: Herbaceous | 0.71 | 0.11 | 6.2 | <0.001 | 0.49 | 0.94 |
| | | Landcover: Open Conifer or Deciduous | 0.27 | 0.05 | 5.8 | <0.001 | 0.18 | 0.36 |
| | | Landcover: Shrub | 0.46 | 0.07 | 7.0 | <0.001 | 0.33 | 0.58 |
| | | logStepLength | 0.08 | 0.00 | 16.3 | <0.001 | 0.07 | 0.09 |
| | | logStepLength:FastState | -0.09 | 0.01 | -16.2 | <0.001 | -0.10 | -0.08 |
| | | Railway | 0.85 | 0.06 | 13.2 | <0.001 | 0.72 | 0.98 |
| | | Slope | -0.01 | 0.07 | -0.1 | 0.887 | -0.14 | 0.12 |

| *Species* | *Season* | *Parameter* | *Estimate* | *SE* | *Wald* | *p-value* | *LCL* | *UCL* |
|---|---|---|---|---|---|---|---|---|
| | | Southerly Aspect | -0.00 | 0.04 | -0.1 | 0.948 | -0.09 | 0.08 |
| | | Trail | 0.11 | 0.14 | 0.8 | 0.449 | -0.17 | 0.39 |
| | | TrailDensity | -0.93 | 0.22 | -4.3 | <0.001 | -1.35 | -0.51 |
| | | TrailDensity:logDistPavedRoad | 0.00 | 0.01 | 0.2 | 0.813 | -0.01 | 0.01 |
| Grizzly Bear | Summer | Burned since 1960 | 0.72 | 0.06 | 12.0 | <0.001 | 0.61 | 0.84 |
| | | Cosine Turn Angle | -1.91 | 0.06 | -31.3 | <0.001 | -2.03 | -1.79 |
| | | Cosine Turn Angle:FastState | 2.66 | 0.07 | 35.8 | <0.001 | 2.51 | 2.80 |
| | | Distance Campground | -0.34 | 0.29 | -1.2 | 0.248 | -0.91 | 0.24 |
| | | Distance to Forest Edge | -2.43 | 0.22 | -10.9 | <0.001 | -2.86 | -1.99 |
| | | Distance to Patch > 9 km2 | -1.33 | 0.05 | -25.6 | <0.001 | -1.43 | -1.23 |
| | | Distance Town | 0.98 | 0.23 | 4.3 | <0.001 | 0.54 | 1.42 |
| | | Distance Town:Night | -0.05 | 0.01 | -3.4 | 0.001 | -0.07 | -0.02 |
| | | Elevation | -0.52 | 0.18 | -2.9 | 0.004 | -0.87 | -0.17 |
| | | Landcover: Barren | 0.71 | 0.19 | 3.7 | <0.001 | 0.34 | 1.07 |
| | | Landcover: Herbaceous | 1.31 | 0.29 | 4.5 | <0.001 | 0.74 | 1.88 |
| | | Landcover: Open Conifer or Deciduous | 0.92 | 0.10 | 9.5 | <0.001 | 0.73 | 1.11 |
| | | Landcover: Shrub | 1.18 | 0.14 | 8.7 | <0.001 | 0.91 | 1.44 |
| | | logStepLength | -1.72 | 0.01 | -116.5 | <0.001 | -1.75 | -1.69 |
| | | logStepLength:FastState | 2.51 | 0.02 | 152.5 | <0.001 | 2.48 | 2.54 |

| *Species* | *Season* | *Parameter* | *Estimate* | *SE* | *Wald* | *p-value* | *LCL* | *UCL* |
|---|---|---|---|---|---|---|---|---|
| | | Railway | -0.42 | 0.19 | -2.2 | 0.027 | -0.80 | -0.05 |
| | | Slope | -0.45 | 0.07 | -6.8 | <0.001 | -0.57 | -0.32 |
| | | Southerly Aspect | 0.14 | 0.06 | 2.4 | 0.017 | 0.02 | 0.25 |
| | | Trail | 0.27 | 0.13 | 2.1 | 0.036 | 0.02 | 0.51 |
| | | TrailDensity | -1.92 | 0.80 | -2.4 | 0.017 | -3.50 | -0.35 |
| | | TrailDensity:logDistPavedRoad | 0.07 | 0.01 | 10.8 | <0.001 | 0.06 | 0.08 |
| Wolf | Fall | Burned since 1960 | 0.03 | 0.05 | 0.5 | 0.592 | -0.07 | 0.12 |
| | | Cosine Turn Angle | -0.83 | 0.04 | -19.1 | <0.001 | -0.91 | -0.74 |
| | | Cosine Turn Angle:FastState | 1.29 | 0.06 | 20.6 | <0.001 | 1.16 | 1.41 |
| | | Distance Campground | 0.15 | 0.39 | 0.4 | 0.695 | -0.62 | 0.93 |
| | | Distance to Forest Edge | -0.30 | 0.13 | -2.4 | 0.016 | -0.55 | -0.06 |
| | | Distance to Patch > 9 km2 | -0.96 | 0.09 | -10.5 | <0.001 | -1.14 | -0.78 |
| | | Distance Town | 1.57 | 0.39 | 4.0 | <0.001 | 0.81 | 2.33 |
| | | Distance Town:Night | 0.03 | 0.02 | 2.0 | 0.048 | 0.00 | 0.06 |
| | | Elevation | -0.75 | 0.20 | -3.8 | <0.001 | -1.14 | -0.36 |
| | | Landcover: Barren | 0.30 | 0.08 | 3.6 | <0.001 | 0.13 | 0.46 |
| | | Landcover: Herbaceous | 0.74 | 0.09 | 8.2 | <0.001 | 0.57 | 0.92 |
| | | Landcover: Open Conifer or Deciduous | 0.34 | 0.06 | 6.2 | <0.001 | 0.24 | 0.45 |
| | | Landcover: Shrub | 0.77 | 0.07 | 11.3 | <0.001 | 0.64 | 0.91 |
| | | logStepLength | -0.27 | 0.01 | -33.0 | <0.001 | -0.28 | -0.25 |
| | | logStepLength:FastState | 0.56 | 0.01 | 61.7 | <0.001 | 0.54 | 0.57 |

| *Species* | *Season* | *Parameter* | *Estimate* | *SE* | *Wald* | *p-value* | *LCL* | *UCL* |
|---|---|---|---|---|---|---|---|---|
| | | Railway | 0.38 | 0.21 | 1.8 | 0.073 | -0.04 | 0.80 |
| | | Slope | -0.77 | 0.06 | -12.7 | <0.001 | -0.89 | -0.65 |
| | | Southerly Aspect | 0.10 | 0.03 | 3.1 | 0.002 | 0.04 | 0.16 |
| | | Trail | 0.18 | 0.13 | 1.4 | 0.163 | -0.07 | 0.44 |
| | | TrailDensity | -0.99 | 0.68 | -1.5 | 0.145 | -2.33 | 0.34 |
| | | TrailDensity:logDistPavedRoad | 0.05 | 0.01 | 4.9 | <0.001 | 0.03 | 0.06 |
| Wolf | Spring | Burned since 1960 | -0.15 | 0.04 | -3.8 | <0.001 | -0.23 | -0.07 |
| | | Cosine Turn Angle | -1.09 | 0.03 | -31.6 | <0.001 | -1.15 | -1.02 |
| | | Cosine Turn Angle:FastState | 1.63 | 0.05 | 32.5 | <0.001 | 1.53 | 1.73 |
| | | Distance Campground | 0.14 | 0.43 | 0.3 | 0.746 | -0.70 | 0.98 |
| | | Distance to Forest Edge | -0.66 | 0.17 | -3.9 | <0.001 | -0.99 | -0.33 |
| | | Distance to Patch > 9 km2 | -0.65 | 0.09 | -7.2 | <0.001 | -0.83 | -0.48 |
| | | Distance Town | 2.08 | 0.48 | 4.4 | <0.001 | 1.15 | 3.02 |
| | | Distance Town:Night | -0.00 | 0.01 | -0.3 | 0.741 | -0.03 | 0.02 |
| | | Elevation | -1.02 | 0.17 | -6.0 | <0.001 | -1.36 | -0.69 |
| | | Landcover: Barren | -0.15 | 0.09 | -1.5 | 0.121 | -0.33 | 0.04 |
| | | Landcover: Herbaceous | 0.33 | 0.09 | 3.6 | <0.001 | 0.15 | 0.51 |
| | | Landcover: Open Conifer or Deciduous | 0.16 | 0.06 | 2.5 | 0.014 | 0.03 | 0.28 |
| | | Landcover: Shrub | 0.52 | 0.08 | 6.9 | <0.001 | 0.38 | 0.67 |

| *Species* | *Season* | *Parameter* | *Estimate* | *SE* | *Wald* | *p-value* | *LCL* | *UCL* |
|---|---|---|---|---|---|---|---|---|
| | | logStepLength | -0.22 | 0.01 | -34.3 | <0.001 | -0.23 | -0.21 |
| | | logStepLength:FastState | 0.46 | 0.01 | 66.6 | <0.001 | 0.45 | 0.48 |
| | | Railway | 0.23 | 0.20 | 1.2 | 0.249 | -0.16 | 0.63 |
| | | Slope | -0.53 | 0.08 | -6.5 | <0.001 | -0.69 | -0.37 |
| | | Southerly Aspect | 0.24 | 0.03 | 6.8 | <0.001 | 0.17 | 0.30 |
| | | Trail | 0.12 | 0.12 | 1.0 | 0.341 | -0.13 | 0.36 |
| | | TrailDensity | 0.15 | 0.45 | 0.3 | 0.735 | -0.72 | 1.03 |
| | | TrailDensity:logDistPavedRoad | 0.02 | 0.01 | 1.8 | 0.075 | -0.00 | 0.04 |
| Wolf | Summer | Burned since 1960 | 0.32 | 0.04 | 7.5 | <0.001 | 0.24 | 0.41 |
| | | Cosine Turn Angle | -0.80 | 0.04 | -20.9 | <0.001 | -0.88 | -0.73 |
| | | Cosine Turn Angle:FastState | 1.24 | 0.05 | 22.8 | <0.001 | 1.13 | 1.35 |
| | | Distance Campground | 1.00 | 0.41 | 2.5 | 0.014 | 0.20 | 1.80 |
| | | Distance to Forest Edge | -0.26 | 0.14 | -1.9 | 0.055 | -0.53 | 0.01 |
| | | Distance to Patch > 9 km2 | -0.63 | 0.05 | -11.7 | <0.001 | -0.74 | -0.53 |
| | | Distance Town | 1.11 | 0.44 | 2.5 | 0.012 | 0.24 | 1.97 |
| | | Distance Town:Night | 0.06 | 0.01 | 4.1 | <0.001 | 0.03 | 0.09 |
| | | Elevation | -0.40 | 0.16 | -2.6 | 0.010 | -0.70 | -0.10 |
| | | Landcover: Barren | 0.15 | 0.13 | 1.1 | 0.268 | -0.11 | 0.41 |
| | | Landcover: Herbaceous | 0.46 | 0.13 | 3.4 | 0.001 | 0.19 | 0.72 |
| | | Landcover: Open Conifer or Deciduous | 0.26 | 0.09 | 3.0 | 0.003 | 0.09 | 0.43 |

| *Species* | *Season* | *Parameter* | *Estimate* | *SE* | *Wald* | *p-value* | *LCL* | *UCL* |
|---|---|---|---|---|---|---|---|---|
| | | Landcover: Shrub | 0.44 | 0.09 | 5.1 | <0.001 | 0.27 | 0.61 |
| | | logStepLength | -0.29 | 0.01 | -40.3 | <0.001 | -0.30 | -0.27 |
| | | logStepLength:FastState | 0.62 | 0.01 | 79.5 | <0.001 | 0.60 | 0.63 |
| | | Railway | 0.23 | 0.23 | 1.0 | 0.311 | -0.21 | 0.67 |
| | | Slope | -1.00 | 0.07 | -14.9 | <0.001 | -1.13 | -0.86 |
| | | Southerly Aspect | 0.18 | 0.04 | 4.0 | <0.001 | 0.09 | 0.26 |
| | | Trail | 0.29 | 0.10 | 3.0 | 0.003 | 0.10 | 0.49 |
| | | TrailDensity | -0.83 | 0.40 | -2.1 | 0.036 | -1.61 | -0.05 |
| | | TrailDensity:logDistPavedRoad | 0.02 | 0.01 | 2.5 | 0.012 | 0.00 | 0.04 |
| Wolf | Winter | Burned since 1960 | 0.11 | 0.02 | 4.4 | <0.001 | 0.06 | 0.15 |
| | | Cosine Turn Angle | -0.87 | 0.02 | -40.7 | <0.001 | -0.91 | -0.83 |
| | | Cosine Turn Angle:FastState | 1.36 | 0.03 | 43.6 | <0.001 | 1.30 | 1.42 |
| | | Distance Campground | -0.32 | 0.19 | -1.7 | 0.092 | -0.68 | 0.05 |
| | | Distance to Forest Edge | -0.52 | 0.11 | -4.9 | <0.001 | -0.73 | -0.31 |
| | | Distance to Patch > 9 km2 | -0.86 | 0.07 | -13.2 | <0.001 | -0.99 | -0.73 |
| | | Distance Town | 1.42 | 0.22 | 6.4 | <0.001 | 0.98 | 1.85 |
| | | Distance Town:Night | -0.02 | 0.01 | -2.3 | 0.022 | -0.04 | -0.00 |
| | | Elevation | -1.12 | 0.10 | -11.0 | <0.001 | -1.32 | -0.92 |
| | | Landcover: Barren | -0.04 | 0.06 | -0.8 | 0.452 | -0.15 | 0.07 |
| | | Landcover: Herbaceous | 0.54 | 0.06 | 9.6 | <0.001 | 0.43 | 0.65 |

| *Species* | *Season* | *Parameter* | *Estimate* | *SE* | *Wald* | *p-value* | *LCL* | *UCL* |
|---|---|---|---|---|---|---|---|---|
| | | Landcover: Open Conifer or Deciduous | 0.15 | 0.04 | 4.0 | <0.001 | 0.08 | 0.22 |
| | | Landcover: Shrub | 0.44 | 0.04 | 11.9 | <0.001 | 0.37 | 0.52 |
| | | logStepLength | -0.28 | 0.00 | -70.8 | <0.001 | -0.29 | -0.28 |
| | | logStepLength:FastState | 0.49 | 0.00 | 108.7 | <0.001 | 0.48 | 0.50 |
| | | Railway | 0.44 | 0.10 | 4.5 | <0.001 | 0.25 | 0.64 |
| | | Slope | -0.43 | 0.04 | -10.6 | <0.001 | -0.51 | -0.35 |
| | | Southerly Aspect | 0.15 | 0.03 | 5.1 | <0.001 | 0.09 | 0.21 |
| | | Trail | 0.21 | 0.08 | 2.6 | 0.009 | 0.05 | 0.36 |
| | | TrailDensity | -0.77 | 0.24 | -3.2 | 0.001 | -1.25 | -0.30 |
| | | TrailDensity:logDistPavedRoad | 0.05 | 0.01 | 9.1 | <0.001 | 0.04 | 0.06 |

### *Section 2.4 Resource selection function results*

Table S4. RSF parameter estimates and 95% confidence intervals for grizzly bears and wolves by season in Banff National Park and surrounding areas, 2002 - 2020.

| *Species* | *Season* | *Parameter* | *Estimate* | *SE* | *Statistic* | *p-value* | *LCL* | *UCL* |
|---|---|---|---|---|---|---|---|---|
| Grizzly Bear | Fall | Burned since 1960 | 1.00 | 0.03 | 36.7 | <0.001 | 0.95 | 1.05 |
| | | Distance Campground | 0.37 | 0.23 | 1.6 | 0.108 | -0.08 | 0.82 |
| | | Distance to Forest Edge | -0.62 | 0.24 | -2.6 | 0.008 | -1.09 | -0.16 |
| | | Distance to Patch > 9 km2 | -1.29 | 0.03 | -49.3 | <0.001 | -1.34 | -1.24 |
| | | Distance Town | -0.81 | 0.13 | -6.1 | <0.001 | -1.07 | -0.55 |
| | | Distance Town:Night | -0.06 | 0.01 | -6.0 | <0.001 | -0.08 | -0.04 |
| | | Elevation | -0.07 | 0.16 | -0.4 | 0.672 | -0.39 | 0.25 |
| | | Landcover: Barren | 0.39 | 0.19 | 2.0 | 0.042 | 0.01 | 0.76 |
| | | Landcover: Herbaceous | 1.19 | 0.17 | 7.0 | <0.001 | 0.86 | 1.52 |
| | | Landcover: Open Conifer or Deciduous | 0.46 | 0.08 | 5.7 | <0.001 | 0.31 | 0.62 |
| | | Landcover: Shrub | 0.81 | 0.16 | 5.0 | <0.001 | 0.49 | 1.13 |
| | | Railway | 2.12 | 0.07 | 28.4 | <0.001 | 1.97 | 2.26 |
| | | Slope | 0.14 | 0.10 | 1.4 | 0.159 | -0.05 | 0.33 |
| | | Southerly Aspect | 0.02 | 0.05 | 0.4 | 0.684 | -0.07 | 0.11 |
| | | Trail | 0.01 | 0.09 | 0.1 | 0.933 | -0.17 | 0.19 |
| | | TrailDensity | -1.11 | 0.53 | -2.1 | 0.036 | -2.15 | -0.07 |
| | | TrailDensity:logDistPavedRoad | 0.01 | 0.00 | 2.8 | 0.005 | 0.00 | 0.02 |

| *Species* | *Season* | *Parameter* | *Estimate* | *SE* | *Statistic* | *p-value* | *LCL* | *UCL* |
|---|---|---|---|---|---|---|---|---|
| Grizzly Bear | Spring | Burned since 1960 | -0.09 | 0.03 | -3.1 | 0.002 | -0.15 | -0.03 |
| | | Distance Campground | 0.45 | 0.11 | 4.2 | <0.001 | 0.24 | 0.66 |
| | | Distance to Forest Edge | -1.50 | 0.14 | -10.8 | <0.001 | -1.77 | -1.23 |
| | | Distance to Patch > 9 km2 | -0.65 | 0.03 | -18.9 | <0.001 | -0.71 | -0.58 |
| | | Distance Town | 0.57 | 0.09 | 6.4 | <0.001 | 0.39 | 0.74 |
| | | Distance Town:Night | -0.02 | 0.01 | -1.9 | 0.053 | -0.04 | 0.00 |
| | | Elevation | -0.94 | 0.16 | -5.9 | <0.001 | -1.26 | -0.63 |
| | | Landcover: Barren | 0.21 | 0.11 | 1.9 | 0.060 | -0.01 | 0.42 |
| | | Landcover: Herbaceous | 0.96 | 0.15 | 6.3 | <0.001 | 0.66 | 1.25 |
| | | Landcover: Open Conifer or Deciduous | 0.45 | 0.06 | 7.3 | <0.001 | 0.33 | 0.57 |
| | | Landcover: Shrub | 0.84 | 0.10 | 8.7 | <0.001 | 0.65 | 1.03 |
| | | Railway | 1.14 | 0.05 | 23.7 | <0.001 | 1.04 | 1.23 |
| | | Slope | 0.01 | 0.09 | 0.1 | 0.927 | -0.17 | 0.19 |
| | | Southerly Aspect | -0.03 | 0.05 | -0.5 | 0.599 | -0.13 | 0.07 |
| | | Trail | 0.07 | 0.16 | 0.4 | 0.669 | -0.24 | 0.37 |
| | | TrailDensity | -1.03 | 0.33 | -3.1 | 0.002 | -1.67 | -0.38 |
| | | TrailDensity:logDistPavedRoad | -0.03 | 0.00 | -9.4 | <0.001 | -0.04 | -0.02 |
| Grizzly Bear | Summer | Burned since 1960 | 0.88 | 0.02 | 40.2 | <0.001 | 0.84 | 0.92 |
| | | Distance Campground | -0.07 | 0.11 | -0.6 | 0.561 | -0.29 | 0.16 |

| *Species* | *Season* | *Parameter* | *Estimate* | *SE* | *Statistic* | *p-value* | *LCL* | *UCL* |
|---|---|---|---|---|---|---|---|---|
| | | Distance to Forest Edge | -1.46 | 0.13 | -11.6 | <0.001 | -1.71 | -1.21 |
| | | Distance to Patch > 9 km2 | -1.06 | 0.03 | -42.0 | <0.001 | -1.11 | -1.01 |
| | | Distance Town | 0.53 | 0.10 | 5.4 | <0.001 | 0.34 | 0.72 |
| | | Distance Town:Night | -0.04 | 0.01 | -4.4 | <0.001 | -0.05 | -0.02 |
| | | Elevation | -0.23 | 0.11 | -2.1 | 0.034 | -0.44 | -0.02 |
| | | Landcover: Barren | 0.51 | 0.11 | 4.8 | <0.001 | 0.30 | 0.72 |
| | | Landcover: Herbaceous | 1.03 | 0.14 | 7.2 | <0.001 | 0.75 | 1.31 |
| | | Landcover: Open Conifer or Deciduous | 0.61 | 0.06 | 9.8 | <0.001 | 0.49 | 0.73 |
| | | Landcover: Shrub | 1.01 | 0.11 | 9.4 | <0.001 | 0.80 | 1.22 |
| | | Railway | 0.02 | 0.11 | 0.1 | 0.881 | -0.19 | 0.23 |
| | | Slope | -0.23 | 0.06 | -3.8 | <0.001 | -0.35 | -0.11 |
| | | Southerly Aspect | 0.07 | 0.05 | 1.4 | 0.163 | -0.03 | 0.17 |
| | | Trail | 0.13 | 0.08 | 1.5 | 0.124 | -0.04 | 0.29 |
| | | TrailDensity | -0.47 | 0.46 | -1.0 | 0.308 | -1.38 | 0.43 |
| | | TrailDensity:logDistPavedRoad | 0.04 | 0.00 | 16.5 | <0.001 | 0.03 | 0.04 |
| Wolf | Fall | Burned since 1960 | 0.22 | 0.03 | 7.6 | <0.001 | 0.16 | 0.28 |
| | | Distance Campground | 0.12 | 0.31 | 0.4 | 0.693 | -0.48 | 0.72 |
| | | Distance to Forest Edge | -0.22 | 0.16 | -1.3 | 0.180 | -0.54 | 0.10 |
| | | Distance to Patch > 9 km2 | -0.79 | 0.06 | -13.9 | <0.001 | -0.90 | -0.68 |
| | | Distance Town | 1.70 | 0.30 | 5.6 | <0.001 | 1.10 | 2.29 |

| *Species* | *Season* | *Parameter* | *Estimate* | *SE* | *Statistic* | *p-value* | *LCL* | *UCL* |
|---|---|---|---|---|---|---|---|---|
| | | Distance Town:Night | 0.01 | 0.01 | 0.8 | 0.400 | -0.01 | 0.04 |
| | | Elevation | -0.83 | 0.18 | -4.6 | <0.001 | -1.18 | -0.48 |
| | | Landcover: Barren | 0.30 | 0.15 | 2.0 | 0.047 | 0.00 | 0.60 |
| | | Landcover: Herbaceous | 0.80 | 0.13 | 6.2 | <0.001 | 0.55 | 1.05 |
| | | Landcover: Open Conifer or Deciduous | 0.30 | 0.07 | 4.4 | <0.001 | 0.17 | 0.43 |
| | | Landcover: Shrub | 0.69 | 0.07 | 10.4 | <0.001 | 0.56 | 0.82 |
| | | Railway | 0.33 | 0.17 | 2.0 | 0.044 | 0.01 | 0.66 |
| | | Slope | -0.92 | 0.10 | -9.6 | <0.001 | -1.10 | -0.73 |
| | | Southerly Aspect | 0.11 | 0.05 | 2.3 | 0.019 | 0.02 | 0.20 |
| | | Trail | 0.17 | 0.10 | 1.8 | 0.075 | -0.02 | 0.36 |
| | | TrailDensity | -0.75 | 0.61 | -1.2 | 0.217 | -1.95 | 0.44 |
| | | TrailDensity:logDistPavedRoad | 0.06 | 0.01 | 11.1 | <0.001 | 0.05 | 0.07 |
| Wolf | Spring | Burned since 1960 | 0.29 | 0.02 | 12.9 | <0.001 | 0.24 | 0.33 |
| | | Distance Campground | 0.57 | 0.31 | 1.8 | 0.068 | -0.04 | 1.19 |
| | | Distance to Forest Edge | -0.75 | 0.16 | -4.7 | <0.001 | -1.07 | -0.44 |
| | | Distance to Patch > 9 km2 | -0.72 | 0.06 | -11.5 | <0.001 | -0.84 | -0.60 |
| | | Distance Town | 2.34 | 0.40 | 5.8 | <0.001 | 1.55 | 3.12 |
| | | Distance Town:Night | 0.00 | 0.01 | 0.2 | 0.810 | -0.02 | 0.02 |
| | | Elevation | -0.98 | 0.17 | -5.9 | <0.001 | -1.31 | -0.66 |
| | | Landcover: Barren | -0.06 | 0.12 | -0.5 | 0.619 | -0.28 | 0.17 |

| *Species* | *Season* | *Parameter* | *Estimate* | *SE* | *Statistic* | *p-value* | *LCL* | *UCL* |
|---|---|---|---|---|---|---|---|---|
| | | Landcover: Herbaceous | 0.49 | 0.12 | 3.9 | <0.001 | 0.25 | 0.74 |
| | | Landcover: Open Conifer or Deciduous | 0.23 | 0.10 | 2.4 | 0.015 | 0.05 | 0.42 |
| | | Landcover: Shrub | 0.80 | 0.14 | 5.6 | <0.001 | 0.52 | 1.08 |
| | | Railway | -0.17 | 0.16 | -1.1 | 0.278 | -0.47 | 0.14 |
| | | Slope | -0.85 | 0.09 | -9.7 | <0.001 | -1.02 | -0.68 |
| | | Southerly Aspect | 0.30 | 0.06 | 5.4 | <0.001 | 0.19 | 0.41 |
| | | Trail | -0.04 | 0.13 | -0.3 | 0.786 | -0.30 | 0.23 |
| | | TrailDensity | 0.33 | 0.48 | 0.7 | 0.485 | -0.60 | 1.26 |
| | | TrailDensity:logDistPavedRoad | 0.02 | 0.01 | 3.1 | 0.002 | 0.01 | 0.03 |
| Wolf | Summer | Burned since 1960 | 0.51 | 0.03 | 19.5 | <0.001 | 0.45 | 0.56 |
| | | Distance Campground | 0.97 | 0.32 | 3.0 | 0.003 | 0.34 | 1.60 |
| | | Distance to Forest Edge | -0.16 | 0.17 | -0.9 | 0.347 | -0.48 | 0.17 |
| | | Distance to Patch > 9 km2 | -0.62 | 0.04 | -17.6 | <0.001 | -0.69 | -0.55 |
| | | Distance Town | 2.57 | 0.41 | 6.2 | <0.001 | 1.76 | 3.38 |
| | | Distance Town:Night | 0.04 | 0.01 | 3.4 | 0.001 | 0.02 | 0.06 |
| | | Elevation | -0.40 | 0.16 | -2.5 | 0.012 | -0.71 | -0.09 |
| | | Landcover: Barren | 0.33 | 0.15 | 2.2 | 0.031 | 0.03 | 0.62 |
| | | Landcover: Herbaceous | 0.63 | 0.17 | 3.8 | <0.001 | 0.31 | 0.96 |
| | | Landcover: Open Conifer or Deciduous | 0.36 | 0.08 | 4.5 | <0.001 | 0.20 | 0.51 |
| | | Landcover: Shrub | 0.65 | 0.10 | 6.3 | <0.001 | 0.45 | 0.86 |
| | | Railway | -0.20 | 0.17 | -1.2 | 0.226 | -0.54 | 0.13 |

| *Species* | *Season* | *Parameter* | *Estimate* | *SE* | *Statistic* | *p-value* | *LCL* | *UCL* |
|---|---|---|---|---|---|---|---|---|
| | | Slope | -1.04 | 0.08 | -12.4 | <0.001 | -1.20 | -0.87 |
| | | Southerly Aspect | 0.19 | 0.05 | 3.9 | <0.001 | 0.10 | 0.29 |
| | | Trail | 0.12 | 0.10 | 1.2 | 0.242 | -0.08 | 0.32 |
| | | TrailDensity | -0.64 | 0.42 | -1.5 | 0.129 | -1.46 | 0.19 |
| | | TrailDensity:logDistPavedRoad | 0.02 | 0.01 | 3.6 | <0.001 | 0.01 | 0.03 |
| Wolf | Winter | Burned since 1960 | 0.24 | 0.01 | 16.9 | <0.001 | 0.21 | 0.27 |
| | | Distance Campground | 0.12 | 0.13 | 0.9 | 0.344 | -0.13 | 0.37 |
| | | Distance to Forest Edge | -0.65 | 0.11 | -6.1 | <0.001 | -0.86 | -0.44 |
| | | Distance to Patch > 9 km2 | -1.26 | 0.05 | -27.1 | <0.001 | -1.35 | -1.17 |
| | | Distance Town | 2.42 | 0.19 | 12.5 | <0.001 | 2.04 | 2.79 |
| | | Distance Town:Night | -0.00 | 0.01 | -0.6 | 0.541 | -0.02 | 0.01 |
| | | Elevation | -1.35 | 0.13 | -10.7 | <0.001 | -1.59 | -1.10 |
| | | Landcover: Barren | -0.13 | 0.08 | -1.6 | 0.112 | -0.29 | 0.03 |
| | | Landcover: Herbaceous | 0.68 | 0.07 | 9.4 | <0.001 | 0.54 | 0.82 |
| | | Landcover: Open Conifer or Deciduous | 0.14 | 0.05 | 2.9 | 0.004 | 0.05 | 0.24 |
| | | Landcover: Shrub | 0.52 | 0.05 | 9.8 | <0.001 | 0.42 | 0.63 |
| | | Railway | 0.17 | 0.08 | 2.2 | 0.028 | 0.02 | 0.32 |
| | | Slope | -0.51 | 0.05 | -10.1 | <0.001 | -0.61 | -0.41 |
| | | Southerly Aspect | 0.24 | 0.04 | 6.8 | <0.001 | 0.17 | 0.31 |
| | | Trail | 0.24 | 0.09 | 2.6 | 0.008 | 0.06 | 0.42 |

| *Species* | *Season* | *Parameter* | *Estimate* | *SE* | *Statistic* | *p-value* | *LCL* | *UCL* |
|---|---|---|---|---|---|---|---|---|
| | | TrailDensity | -0.74 | 0.26 | -2.8 | 0.005 | -1.25 | -0.22 |
| | | TrailDensity:logDistPavedRoad | 0.05 | 0.00 | 15.0 | <0.001 | 0.04 | 0.06 |
